# Evolution under Domestication: Genetic differentiation in black soldier fly (*Hermetia illucens*) populations subjected to recent selective breeding

**DOI:** 10.1101/2025.04.14.648847

**Authors:** Shaktheeshwari Silvaraju, Rebecca Loh Ker, Sandra Kittelmann, Nalini Puniamoorthy

## Abstract

The black soldier fly (BSF; *Hermetia illucens*) is widely utilized in commercial and research applications for waste bioconversion and sustainable protein production. However, prolonged captivity and artificial selection can shape genetic diversity, potentially influencing adaptability and long-term population stability. This study examined how recent selective breeding, genetic drift, and relaxed selection have influenced genetic differentiation in BSF populations over a short timeframe. Using mitochondrial cytochrome oxidase I (*CO1*) and genome-wide RAD sequencing, population structure, heterozygosity, and selection signatures across eleven BSF populations, including selectively bred, wild-derived, and commercial strains were analysed. Results revealed that rapid genetic shifts have occurred within selectively bred populations (LA to LE) over ∼5 years, driven by artificial selection, subsequent relaxation, and environmental adaptation. The decline in effective population size (Ne) observed post-COVID-19 suggests recent bottlenecks, which may have further contributed to genetic drift and differentiation. While domesticated populations exhibited reduced genetic diversity and signs of inbreeding, wild-type population under short captivity retained higher heterozygosity. Genome-wide analyses further identified adaptive divergence among populations, with balancing selection and selective sweeps shaping genetic variation. Notably, despite shared ancestry, genetic differentiation persisted in selectively bred populations, reinforcing that selection and environmental pressures continue to influence their genomic landscape even after targeted selection was relaxed. These findings underscore the need to monitor genetic diversity in BSF breeding programs to maintain adaptability, enhance resilience, and mitigate risks from artificial selection and population collapse.

## Introduction

In recent years, research and commercialisation of black soldier flies (BSFs) have advanced substantially with BSF production facilities established all over the world; from South Africa to Spain, The Netherlands, USA (Kentucky), Chile, Malaysia, Singapore, France, Ireland, and many more (Gabriel Patrick, 2021). This peak in the interest in BSFs, driven by the need to develop sustainable protein alternatives, has resulted in numerous studies emerging with regards to improving their performance (Bekker et al., 2021; Gebiola et al., 2023) and body composition (Shelomi, 2020), optimizing their rearing conditions (Sheppard et al., 2002; Spranghers et al., 2017; Yakti et al., 2022), understanding their nutritional needs (Barragan-Fonseca et al., 2019; Chen et al., 2023; Zhang et al., 2023). The gut microbiota is believed to play an important role and thus has been studied frequently in relation to these parameters (Jiang et al., 2019; Klammsteiner et al., 2020; Raimondi et al., 2020; Silvaraju et al., 2024; Yang et al., 2021; Zhineng et al., 2021). Gut microbiota, for instance, has shown to play a crucial role in BSF development, digestion, and metabolism, influencing nutrient absorption and waste degradation efficiency, making them essential for larval performance (Eke et al., 2023). However, despite maintaining similar environmental conditions, different studies reported similarities as well as differences in the core gut microbiota. For instance, even though chicken feed was used as a control diet, Klammsteiner et al., (2020) reported *Actinomyces, Actinomycetales* unclassified, *Dysgonomonas, Enterococcus* as the core gut microbiota present in the BSF larvae irrespective of changing diets (Klammsteiner et al., 2020) while Vandeweyer et al., (2023) reported *Buttiauxella, Enterococcus, Providencia*, and *Morganella* in their study (Vandeweyer et al., 2023). These contradictions have raised questions regarding the influence of genetics on the phenotypic responses (e.g., gut microbiota composition) to environmental variations (eg., diet) (Sandrock et al., 2022). To disentangle the web of mechanisms underlying BSF responses, it is essential first to determine the genetic diversity across populations. To assess this, several methods have been employed thus far.

Over the past decade, studies have investigated the population genetics of BSFs using maternally inherited mitochondrial cytochrome oxidase I (*CO1*) and higher-resolution microsatellite markers to obtain a deeper understanding of their evolutionary trajectory across the world. These studies have revealed that many of the populations present in Europe, Africa, and Asia were introduced from the Americas (Ståhls et al., 2020) and most domesticated populations can be traced back to a source population in North America (Kaya et al., 2021). Although unique genetic profiles were found in certain wild and domesticated populations, domesticated populations across the world appeared to have significantly reduced genetic diversity overall (Guilliet et al., 2022; Kaya et al., 2021), highlighting the dangers of currently uncontrolled human-mediated dispersal events (Athanassiou et al., 2024; Guilliet et al., 2022; Kaya et al., 2021), domesticated populations with small effective size, intensive inbreeding (L. Hoffmann et al., 2021), and even reproductive isolation in certain regions (Park et al., 2017).

The implications of genetic variation in BSF populations even extends to observable morphological differences, including variations in body and abdominal window size, wing and abdominal coloration, and the proportion of white and black pigmentation on the heads (Ståhls et al., 2020). These genetic differences also manifest as variations in performance and body composition such as pupation rate, harvesting weight of pupae, adaptability to various diets (ie., gut microbiota composition), and protein and lipid content of larvae just to name a few (Athanassiou et al., 2024; Greenwood et al., 2021; Khamis et al., 2020; Sandrock et al., 2022; Silvaraju et al., 2024; Zhang et al., 2023).

In a rapidly growing industry focused on BSF commercialization and research, it is essential to carefully characterize the genetic structure of populations to ensure reproducibility. Prolonged captivity impacts the genetic structure of populations, including balancing selection on genes primarily associated with metabolism and developmental processes, or selective sweeps that result in a loss of genetic diversity (Athanassiou et al., 2024; Generalovic et al., 2021). Such a reduction in genetic diversity critically limits the potential of selective breeding programs, where diversity is crucial for the successful selection of traits, and prevention of population collapse under stress (L. Hoffmann et al., 2021; Rhode et al., 2020). However, achieving this clarity is often challenging due to colonies being transferred between facilities without proper documentation of their origins (Rhode et al., 2020).

This study aimed to assess the genetic structure, diversity, and evolutionary relationships of eleven BSF populations including five selectively bred lines (LA to LE) derived from an admixed founder population, three wild-caught populations (WT, SWT, and KR) of which WT and SWT were subsequently established in the lab, one long-term domesticated population (SWS), and two commercial populations (IFT and NT). To achieve this, a dual approach was employed: 1) Partial *CO1* sequences were obtained and compared with published sequences of BSFs from all over the world, to contextualize the evolutionary history of the studied populations within a broader framework, followed by an isolated analysis of the studied populations to provide insights into maternal lineage diversity and phylogenetic relationships; and 2) Restriction site-associated DNA (RAD) sequencing to generate genome-wide high-resolution data, enabling the assessment of population genetic structure, levels of heterozygosity, inbreeding, fine-scale genomic differences across populations, and potential genomic regions under selection. Together, these analyses would provide a foundation for future research on these populations, e.g., the correlation of genetics with gut microbiota composition and larval performance.

## Methods

### Sample Collection and DNA extraction

Black soldier fly (BSF) colonies used in this study were primarily housed at the National University of Singapore (NUS) BSF facility (Table 1). Populations LA to LE were established in 2018 by crossing Southeast Asian and North American cultures (Tok Wei Xian Eugene, 2019), KR, WT and SWT were wild-caught populations from 2018, 2020, and 2022 respectively from which WT, and SWT were established in the facility thereafter. SWS is a domesticated population that originated from the Swiss Federal Institute of Aquatic Science and Technology (Eawag) and was later established in NUS facility in 2023. IFT and NT are populations from Singapore and Malaysia BSF commercial production facilities respectively.

**Table 1.** BSF populations used in the study with corresponding information on the country of collection from (“Collection country”), the country the population was originally sourced from (“Country of origin”), captivity status, and the number of individuals obtained for each population that was utilised in this study.

| Population | Collection country | Country of origin | Status | Sample size (n) |
| --- | --- | --- | --- | --- |
| LA | Singapore | KR (Singapore) x<br>Malaysia x<br>USA x<br>Spain | Domesticated | 20 |
| LB | Singapore |  | Domesticated | 20 |
| LC | Singapore |  | Domesticated | 20 |
| LD | Singapore |  | Domesticated | 20 |
| LE | Singapore |  | Domesticated | 20 |
| WT | Singapore | Central-southern,<br>Singapore | Wild | 20 |
| SWS | Singapore | Switzerland | Domesticated | 20 |
| SWT | Singapore | North-eastern,<br>Singapore | Wild | 20 |
| KR | Singapore | South-western,<br>Singapore | Wild | 5 |
| IFT | Singapore | Singapore | Domesticated | 15 |
| NT | Malaysia | Malaysia | Domesticated | 15 |

Upon collection, larvae were frozen at –80°C until dissection. Whole larval heads were excised on a sterile surface using a sterile scalpel (Kaya et al., 2021). Genomic DNA was extracted from the dissected head of each individual using the QIAamp^®^ PowerFecal^®^ Pro DNA Kit (QIAGEN) following the manufacturer’s handbook, employing the vortex adapter method for the homogenization of tissue samples. Before the final centrifugation step, 50 µl of C6 solution was added onto the MB Spin Column filter membrane and left to incubate at room temperature for 5 min to ensure sufficient time was provided for the DNA to completely dislodge from the membrane.

Extracted DNA was treated with RNase A to remove any residual RNA. The quality and concentration of DNA were verified using 1% agarose gel and Quantus fluorometer to ensure high-quality DNA suitable for sequencing.

### Mitochondrial cytochrome oxidase I (*CO1*) gene amplification and sequencing

The *CO1* gene was amplified from BSF extracted genomic DNA via polymerase chain reaction (PCR) amplification using universal animal (or metazoan) primers mICO1intF (5′-GGWACWGGWTGAACWGTWTAYCCYCC) (Leray et al., 2013) and jgHCO2198 (5′-TAIACYTCIGGRTGICCRAARAAYCA) (Geller et al., 2013), targeting the 313bp *CO1* minibarcode. All primers had unique 13-bp sequences attached to their 5’-ends, which allowed for the simultaneous sequencing of numerous samples on Next Generation Sequencing platform (also known as multiplexing). Primer tags published by Srivathsan and colleagues (2018, 2021) were adopted in this method. Multiplexing was done for each DNA sequencing run, by using a unique combination of forward and reverse primer tags for every sample. Each PCR reaction contained, 5 μL of Rapid Taq Master Mix (Vazyme, China), 1 μL of each primer (5 μM), and 1 μL of 3x-diluted genomic DNA. The PCR cycling conditions were as follows: 3 min initial denaturation at 94°C, followed by 40 cycles of denaturation at 94°C (0.5), annealing at 45°C (1 min), extension at 72°C (0.5 min), and a final extension step at 72°C (3 min). PCR success was confirmed via gel electrophoresis, and PCR products were pooled in equal volumes. The pool of PCR products was purified using the ExoSAP-IT Express PCR product cleanup (Applied Biosystems, USA) followed by Ampure XP bead-based purification (Beckman Coulter, USA), in accordance with the manufacturer’s instructions.

The purified pool was processed for library preparation and DNA sequencing on the MinION Mk1B instrument (Oxford Nanopore Technologies, UK). Library preparation was done using the SQK-LSK114 Ligation Sequencing Kit and sequenced on the R10.4 flow cell, generating output files in the POD format. Base-calling of the POD files was performed using Dorado version 0.4.0 (Oxford Nanopore Technologies) with the duplex base-calling option enabled. The resulting FASTQ files were demultiplexed and underwent quality control processing using ONTbarcoder version 0.1.9 (Srivathsan et al., 2021) with default settings, except for two modifications: barcode length was set to 313 bp to accommodate potential *CO1* sequence variation, allowing up to four codons (12 bp) to be missing, and barcode minimum length was adjusted to 300 bp (excluding primers and indices).

### Sequence assembly and alignment

In addition to the samples from above, *CO1* sequences of BSF global populations available were downloaded and collated for comparative analysis (Ebeneezar et al., 2021; Guilliet et al., 2022; Park et al., 2017; Sandrock et al., 2022; Ståhls et al., 2020; number of sequences and their GenBank Accession Numbers provided in Table S1). All sequences (n = 329) were processed using Molecular Evolutionary Genetic Analysis (MEGA) software version 10.2.4. Sequence alignment was conducted using MUSCLE with default settings. Low-quality sequences with too many ambiguous bases were removed (8 sequences removed). The remaining 516 sequences were trimmed by 353 bases from the 5’ end to match those obtained from the populations used in this study. Additionally, the last base (site 667) at the 3’ end was removed to eliminate an ambiguous base present in publicly available sequences. The final length of all aligned sequences was 314 bases. The aligned sequences were saved in FASTA format and imported into R for population genetic and phylogenetic analysis.

### *CO1* data bioinformatic processing

Neighbour-Joining (NJ) tree was constructed to infer phylogenetic relationships among BSF populations using ape (version 5.7-1). Genetic distances were calculated using the Prevosti distance metric with 100 bootstraps performed to assess node support, with only bootstrap values above 90% displayed on the tree. Genetic diversity and differentiation analyses were conducted by calculating overall haplotype diversity followed by diversity per population (pegas; version 1.3). Nucleotide diversity (π) was calculated for overall populations and per population, and pairwise FST values were computed using hierfstat (version 0.5-11) and summarized in a heatmap to visualize genetic differentiation between populations. Mean gene diversity (Hs) within populations was also calculated to provide insights into within-population genetic variation. Analysis of molecular variance (AMOVA) was conducted on the genetic distance matrix generated from the *CO1* sequences to partition genetic variance within and among populations (pegas; version 1.3). Haplogroups were identified in populations used in this study as well as published global population data (haplotypes package; version 1.1.3.1) and visualised as pie charts using ggplot2. Pie charts were then manually inserted onto a world map for informative visualisation of global genetic lineages.

### Restriction site-associated DNA sequencing (RAD-seq)

Genomic DNA was sent to SNPsaurus (Oregon, United States) for the preparation of nextRAD libraries targeting approximately 15,000 loci. The nextRAD library preparation method involves fragmentation of genomic DNA using Nextera reagents, followed by selective amplification with primers that include adapter sequences and extend into genomic DNA with selective bases at the 3’ end, allowing consistent enrichment of a defined subset of loci across samples. Library preparation and sequencing were conducted on an Illumina platform, generating single-end 122 bp reads with an average read depth of 20× per locus. Raw sequencing data in BCL format were demultiplexed and converted to FASTQ files using bcl-convert (v4.2.4; Illumina). Reads were then quality-filtered and aligned to the *Hermetia illucens* reference genome (GCF_905115235.1_iHerIll2.2.curated.20191125_genomic.fna). Single Nucleotide Polymorphism (SNP) calling was performed using SNPsaurus’ custom variant-calling pipeline. The resulting variant calls (SNPs) were provided in VCF format and imported into RStudio (version 2024.09.1) for downstream population genomic analysis.

### RAD-seq bioinformatic analysis Principal Components Analysis

The VCF file was imported and converted into a genlight object using the vcfR (version 1.15.0) and adegenet (version 2.1.10) packages for downstream population genetic analyses. To assess genetic differentiation among populations, a Principal Components Analysis (PCA) was conducted using the glPca function from the adegenet package PCA was performed with three retained components (nf = 3), and the variance explained by each component was calculated and visualized. The results were plotted using ggplot2 (version 3.5.1).

### Phylogenetic Analysis

Phylogenetic relationships among individuals and populations were assessed using UPGMA (Unweighted Pair Group Method with Arithmetic Mean) clustering. Genetic distance matrices were calculated using the bitwise.dist function (individual-level) and prevosti.dist (population-level) from the poppr package. Phylogenetic trees with bootstrap support (100 replicates) were generated using the aboot function and visualized with plot.phylo from the ape package (version 5.7-1).

### Genetic Diversity and Differentiation Metrics

Genetic diversity indices including observed (Ho) and expected heterozygosity (He) and inbreeding coefficients (FIS) were calculated using the basic.stats function from the hierfstat package (version 0.5-11). Pairwise genetic differentiation (Fst) between populations was estimated using Weir and Cockerham’s (1984) method implemented in the genet.dist function. Results were visualized in heatmap format with annotated Fst values using ggplot2.

### Analysis of Molecular Variance

Analysis of Molecular Variance (AMOVA) was conducted using the poppr.amova function from the poppr package (version 2.9.3). The hierarchical structure was specified with populations defined by metadata. Permutation tests (999 replicates) were performed to assess the significance of variance components.

### Admixture Analysis

Genotype data were first converted into STRUCTURE format using a custom R function following Tom Jenkins’ tutorial (https://github.com/Tom-Jenkins/admixture_pie_chart_map_tutorial/blob/master/TJ_genind2structure_function.R), and subsequently used for admixture analysis. Admixture proportions were estimated using sparse non-negative matrix factorization (sNMF) with the snmf function from the LEA package (version 3.6.0). A range of K values (1–10) was tested with 10 repetitions per K. The optimal number of clusters (K = 6) was determined by selecting the K value with the lowest cross-entropy value across replicates. The best Q-matrix was extracted, and individual ancestry coefficients were visualized using ggplot2. Samples were grouped by predefined populations.

### Tajima’s D Calculation and Annotation

To identify signatures of selection, Tajima’s D was calculated for each population using vcftools (version 0.1.16) on Ubuntu Linux with a sliding window approach (window size = 50 kb; step size = 10 kb). Resulting output files were converted into BED format and annotated using bedtools (version 2.30.0) against *Hermetia illucens* genome (NCBI; GCF_905115235.1_iHerIll2.2.curated.20191125_genomic.fna). Annotated windows were imported into RStudio for genome-wide visualization of Tajima’s D values using ggplot2. Candidate genomic regions under selection were identified by examining outlier windows with extreme Tajima’s D values and cross-referencing these regions with annotated genes to infer potential functional relevance.

### Demographic Inference via Stairway Plot

For demographic inference, populations LA–LE were selected based on metadata that assigned each individual (AccessID) to its respective population. Site frequency spectra (SFS) were computed for each subset population using the pegas package (version 1.3), with the SFS folded due to the absence of an ancestral reference. Input blueprints for each population included parameters such as sample size, total observed nucleotide sites (L), mutation rate (3.5 × 10^−9^ per site per generation, based on *Drosophila* as a proxy) (Keightley et al., 2009), generation time (40 days per generation) (Slagboom et al., 2024), and the timeframe of 2018 – 2024. Stairway Plot was executed via Java batch scripts to estimate historical effective population sizes (Ne). Final outputs were compiled and visualized in RStudio using ggplot2 to depict changes in Ne over time.

## Results

### Population structure of recently selectively bred populations based on *CO1*

Mitochondrial *COI-*based genetic analysis was performed on an average of 4,158 ± 2,260 sequences assigned to each sample whereby all sequences were of high quality and none of them had ambiguous bases after post-processing using ONTbarcoder. A total of 194 individuals, 23 loci, and 46 alleles were analysed from populations used in this study.

Differences in population structure across the selectively bred, wild-derived, and commercial populations used in this study were visualized based on maternal haplogroups, defined by nucleotide differences between samples (Figure 1). The populations clustered into five haplogroups. Based on maternal haplotype diversity (Hd) and nucleotide diversity (π) metrics, the overall haplotype diversity across all populations was relatively low (0.296), while a moderate nucleotide diversity (0.009) was observed, indicating that the few haplotypes present were highly diverse.

**Figure 1.**
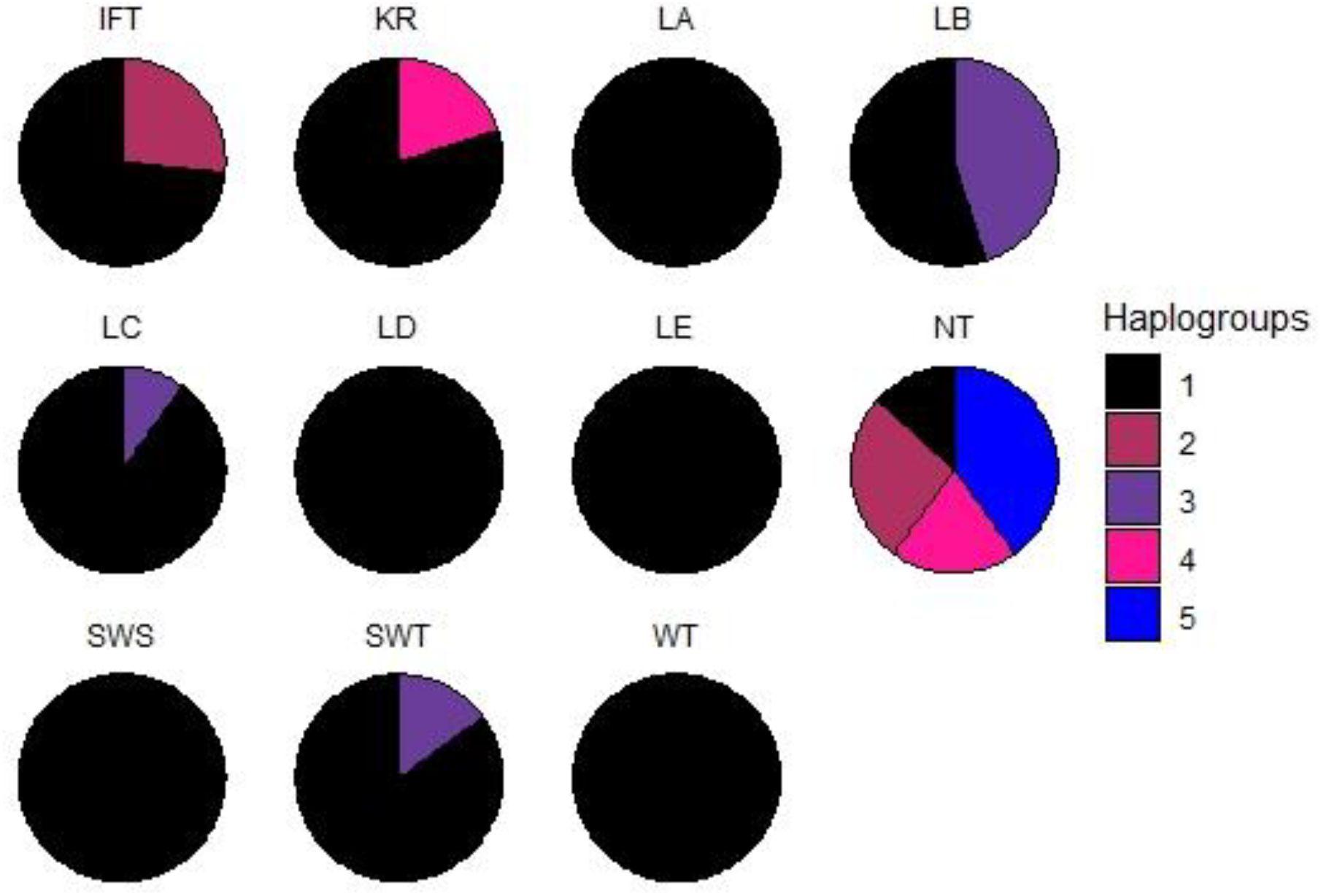
Proportional distribution of mitochondrial haplogroups across domesticated and wild BSF populations. Each pie chart represents a population and is divided into segments corresponding to haplogroup frequencies. The colours represent different haplogroups (1–5) as shown in the legend. Refer to Table 1 for details of population abbreviations.

Significant genetic variation was observed among BSF populations, with some populations (LA, LD, LE, WT, and SWS) exhibiting no genetic diversity (Hd = 0, π = 0), dominated by a single haplogroup, suggesting potential bottleneck events or inbreeding (Table S2; Figure 1). In contrast, LB and LC displayed moderate to low haplotype diversity (Hd = 0.521 and 0.189, respectively) with low nucleotide diversity, despite originating from the same admixed population as LA, LD, and LE. Notably, the commercial populations IFT and NT exhibited the highest nucleotide diversity (π > 0.02), with NT showing the greatest haplotype diversity (Hd = 0.762), reflecting substantial gene flow and a larger effective population size.

The Neighbour-Joining tree (Figure S1) further supported these genetic relationships, revealing relatively limited genetic distances among most populations (Fst < 0.05; Figure S2A). Despite their shared admixed origins, LA was genetically closer to the wild-caught WT than to LB or LC, while LB showed greater genetic similarity to SWT. In contrast, KR, IFT, and NT formed distinct branches, with NT exhibiting the highest Fst values (≥ 0.1), indicating a greater degree of genetic differentiation. However, the largest observed genetic distance was 0.6, with an Fst value of 0.146, suggesting that while some divergence exists, the populations share a relatively recent common ancestry or ongoing gene flow.

To assess how much of the variance observed was due to genetic differentiation between populations and within individual populations, Analysis of Molecular Variance (AMOVA) was performed. Approximately 57.04% of the total genetic variation was attributable to differences between populations, while the remaining 42.96% of the variation occurred within populations (PhiST = 0.57, *p* < 0.001), indicating genetic differentiation among populations to an extent.

### Population structure based on *CO1* sequences

To contextualize the studied populations within a broader global structure, publicly available global *CO1* datasets of BSF populations (Materials and Methods section; Table S1) from different regions were obtained and utilized as a proxy. After filtering low-quality sequences and any ambiguous bases, the final length of all aligned sequences was 314 bases, with no ambiguous bases remaining. A total of 516 sequences from 58 geographic regions, 29 countries, and 6 continents, inclusive of data from the current study, were analysed. The number of sequences per country ranged from 1 to 244, with South Korea contributing the largest dataset.

Global genetic relationships among BSF populations were visualized using a Neighbour-Joining phylogenetic tree (Figure S3A), revealing distinct clustering based on geographic origins. Asian populations predominantly formed well-supported clades, while those from Africa, Europe, Australia, South America, and North America were more dispersed.

Population differences were further examined through maternal haplogroups (Figure 2). Notably, African and Central American populations exhibited highly homogeneous haplogroup distributions despite close proximity, suggesting limited gene flow. In contrast, South American populations showed admixture, with shared haplogroups between Peru and Paraguay (haplogroup 6), and Bolivia and Brazil (haplogroup 5). Bolivia exhibited the highest haplogroup diversity, containing five unique haplogroups (19–23) absent elsewhere. Certain countries harboured unique haplogroups, potentially due to localised adaptation for example Australia (haplogroup 9), Guyana (haplogroup 13), Brazil (haplogroups 17 & 18), and South Korea (haplogroup 3).

**Figure 2.**
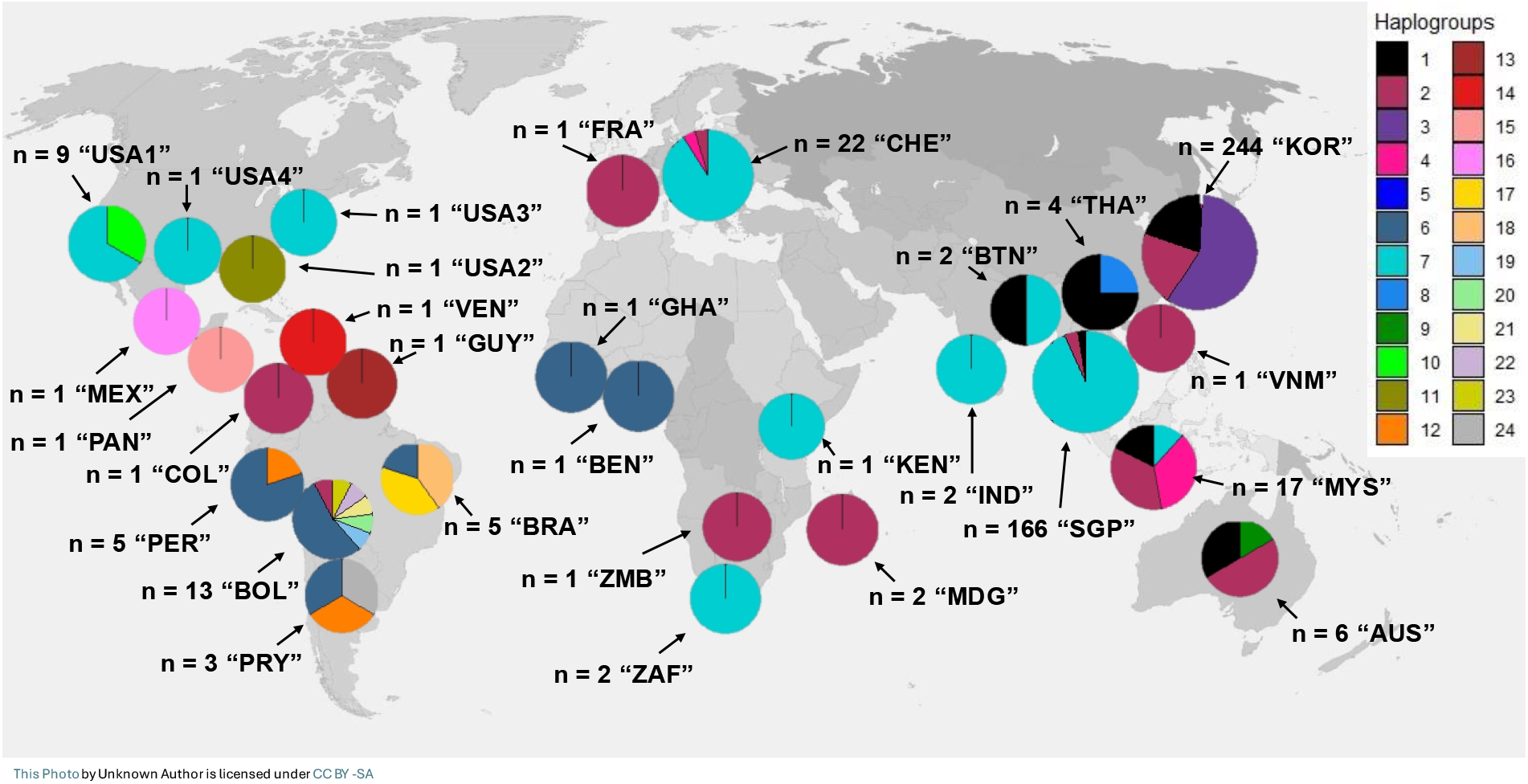
Global distribution of BSF haplogroups based on publicly available *CO1* data. Each pie chart represents a specific country and is divided into segments corresponding to haplogroup frequencies. The colours represent different haplogroups (1–24) as shown in the legend, which were assigned based on a hierarchical clustering algorithm applied to genetic distance data (h = 2) grouping samples with up to 2 base pair (bp) differences into the same haplogroup. Pie chart sizes are proportional to sample sizes, with larger pies representing larger sample sizes and smaller pies indicating fewer individuals. Pie charts representing individual regions are provided in Figure S3B. Countries included in the dataset are AUS: Australia (6), BEN: Benin (1), BTN: Bhutan (2), BOL: Bolivia (13), BRA: Brazil (5), USA1: California (9), COL: Colombia (1), USA2: Florida (1), FRA: France (1), GHA: Ghana (1), GUY: Guyana (1), KEN: Kenya (1), IND: Kerala (2), MDG: Madagascar (2), MYS: Malaysia (17), MEX: Mexico (1), USA3: North Carolina (1), PAN: Panama (1), PRY: Paraguay (3), PER: Peru (5), SGP: Singapore (166), ZAF: South Africa (2), KOR: South Korea (244), CHE: Switzerland (22), USA4: Texas (1), THA: Thailand (4), VEN: Venezuela (1), VNM: Vietnam (1), ZMB: Zambia (1), with the number of sequences included in brackets.

Notably, the Singapore BSF populations used in this study were dominated by haplogroup 7 which was also found in populations from India, Africa, Switzerland, California, and Texas (Figure S3). A smaller portion belonged to haplogroup 2, shared with populations from Australia, Colombia, France, Madagascar, Malaysia, Vietnam, Zambia, and South Korea, reflecting the geographic dominance of haplogroups 2 and 7 across multiple continents.

While mitochondrial data provided valuable insights into the characterization of the studied populations within the global *CO1* structure and broader patterns of maternal variation, it may not have fully captured the genetic complexity of these populations. To achieve a more comprehensive understanding, a genome-wide analysis using RAD sequencing was conducted.

### Population structure and genetic variance based on RAD-seq

RAD-seq was performed on all populations, both domesticated and wild, used in this study to obtain a further in-depth view of the population structure. An average of 4,724,995 ± 693,911 high-quality reads were obtained with an average of 3363251 ± 715720 (∼70%) of these reads mapped to the reference genome. The genotyping data consisted of individuals with diploid genomes (ploidy = 2), represented by 22,813 SNP loci distributed across chromosomes, assigned to respective populations.

Genetic structure among BSF populations used in this study was assessed using Principal Components Analysis (PCA; Figure 3). The PCA plot revealed clear genetic structure among populations, with some forming distinct clusters while others exhibited overlaps. SWS and WT individually formed two distinct clusters that were spatially separated from all other populations, suggesting significant genetic differences. In contrast, NT and KR clustered together, while populations LA, LD, and LE exhibited considerable overlap. Similarly, LB, LC, and SWT also formed an overlapping cluster, indicating relatively low genome-wide differentiation among these groups. AMOVA based on high-resolution, genome-wide nuclear DNA revealed that the majority of genetic variation occurred within populations (72.17%), while only 27.82% was attributed to differences between populations (PhiST = 0.278, *p* = 0.01). This confirmed that while some populations exhibited distinct clustering, genetic variation was largely shared within populations.

**Figure 3.**
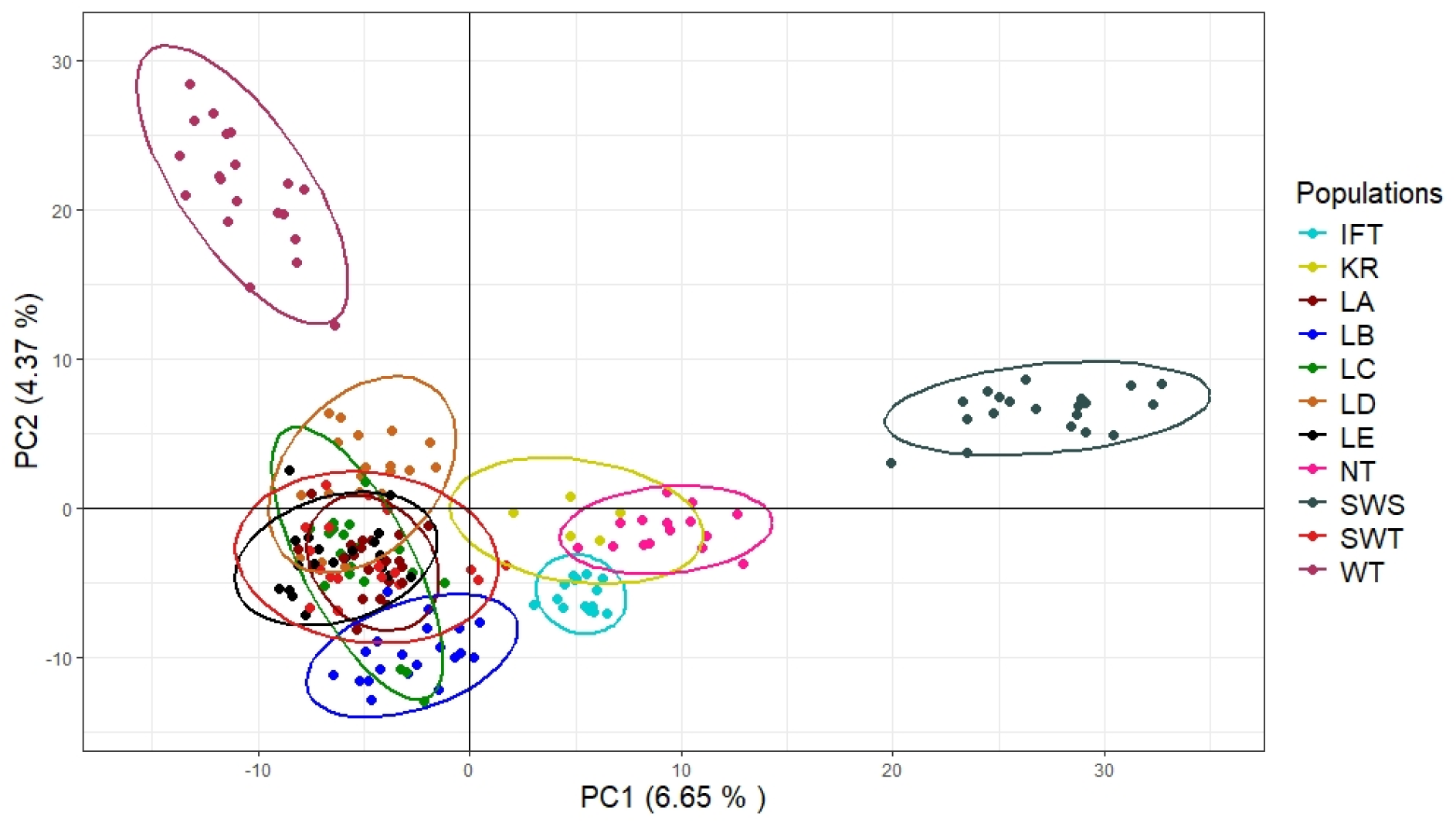
Principal Components Analysis (PCA) plot illustrating the genetic structure among BSF populations. Each colour represents a distinct population, with individual points corresponding to individual samples within each population. Ellipses indicate the 95% confidence intervals for each population.

To further examine population structure at a finer scale, admixture analysis using sparse non-negative matrix factorization (sNMF) was conducted (Figure 4). This approach estimated the proportion of ancestry in each population, identifying six inferred genetic clusters (K = 6). The analysis showed considerable variation in ancestry proportions among individuals; notably, populations LC (Cluster 4), LD (Cluster 6), LE (Cluster 2), WT (Cluster 1), SWT (Cluster 1), KR (Cluster 3), and IFT (primarily Cluster 4) were each dominated by a single ancestral cluster, indicating relatively little admixture. In contrast, LA, LB, SWS, and NT exhibited substantial contributions from multiple ancestral clusters. Interestingly, despite originating from the same admixed population, populations LA to LE have diverged significantly in their admixture proportions, indicating substantial genetic differentiation over a short period of time (∼5 years). While PCA analysis indicated broad genetic similarity among LA to LE, the divergence observed in these populations suggested that selective breeding and/or environmental adaptation had actively driven differentiation, leading to recent and ongoing rapid genetic shifts even within populations of shared ancestry, despite the absence of large-scale genomic divergence.

**Figure 4.**
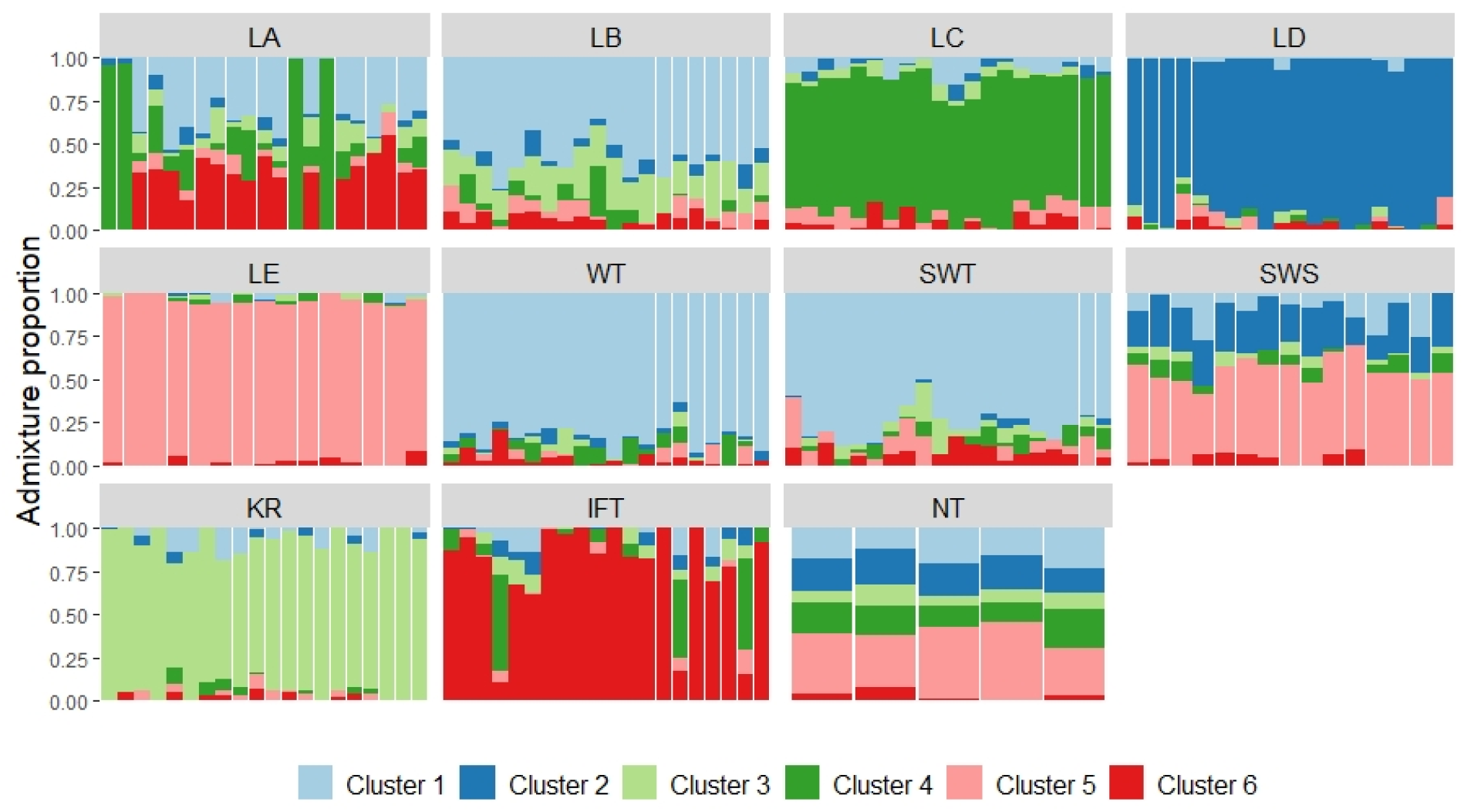
Admixture proportions across the studied BSF populations for K = 6 inferred genetic clusters. Each panel represents a predefined population, with each vertical bar corresponding to an individual sample. Colours represent the six inferred clusters (Cluster 1–6) and the proportion of genetic ancestry each individual derived from. The y-axis shows admixture proportions (0–1). Certain populations were dominated by a single cluster (e.g., Cluster 5 in LC, Cluster 6 in LD, Cluster 3 in KR, Cluster 2 in LE, and Cluster 1 in WT and SWT), while others (LA, LB, SWS, NT) exhibited substantial admixture from multiple clusters.

While admixture analysis revealed the degree of genetic mixing across populations, further insights into the genetic structure (ie., within-population diversity and between-population diversity) was obtained through the calculation of Fis (inbreeding coefficient) to assess the level of inbreeding or deviation from Hardy-Weinberg equilibrium within each population, and Fst (fixation index; Figure S2B) to quantify genetic differentiation between populations.

Overall, the Fis values ranged from 0.121 to 0.234, indicating varying levels of inbreeding across populations. The highest values were observed in IFT (0.234), NT (0.227), and LB (0.226), suggesting a greater degree of inbreeding or heterozygosity deficit compared to expectations (Figure S4). Notably, IFT and NT are commercial populations, while LB is an inbred population, potentially explaining their elevated Fis values. In contrast, the wild-caught populations, WT (0.121) and SWT (0.144) exhibited the lowest Fis values, indicating relatively lower levels of inbreeding. Most other populations, including LA (0.190), LC (0.204), LD (0.224), LE (0.194), and SWS (0.222), displayed moderate Fis values, potentially reflecting restricted gene flow in captivity or selective breeding practices.

When genetic differentiation between populations was analysed (Figure S2B) SWS and WT were observed to be highly differentiated (Fst = 0.373), followed by SWS with all the other populations, and WT with all the other populations (Fst range 0.183 to 0.293). This supports the hypothesis that their wild-caught status (WT) and geographic origin (SWS) contributed to their unique genetic profiles. Conversely, populations other than WT and SWS exhibited relatively low Fst values, indicating minimal differences in allele frequencies among them. The low genetic differentiation among domesticated populations, despite observable shifts in admixture proportions, suggested that while selection and adaptation were influencing genetic composition, genome-wide divergence remained limited at this stage.

### Genetic effects of domestication

To explore the demographic processes underlying the genetic differentiation and shifts in genetic composition observed in LA to LE, effective population size (Ne) trajectories of the selectively bred populations were reconstructed using a Stairway Plot (Figure 5), capturing the effects of artificial selection and subsequent relaxed selection that took place from 2018 to 2024.

**Figure 5.**
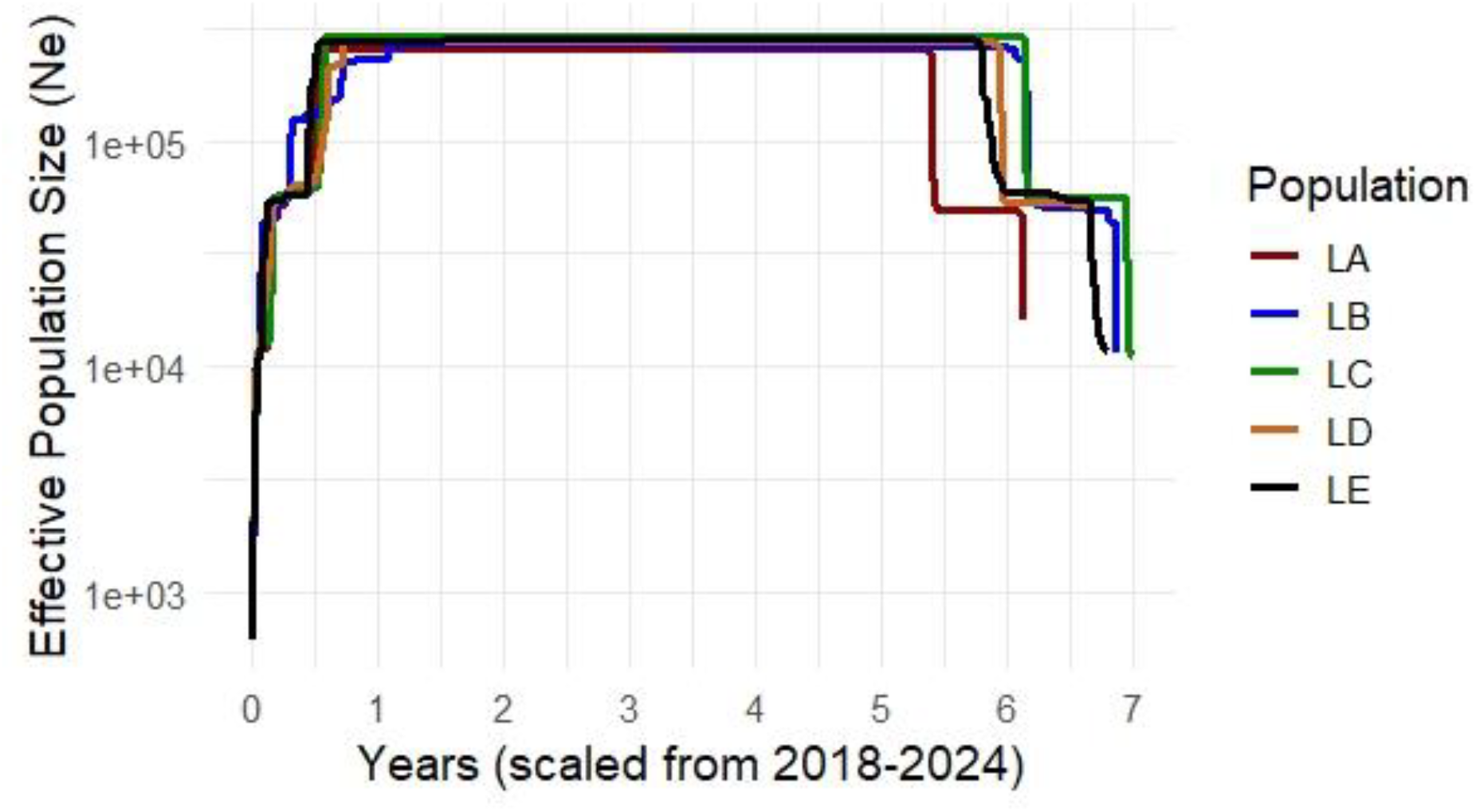
Stairway plot displaying the demographic history of selectively bred BSF populations (LA to LE) from 2018 to 2024. The x-axis represents time (years) scaled from 2018 to 2024, while the y-axis represents the effective population size (Ne) on a log scale.

The demographic analysis revealed a sharp increase in Ne during the initial phase of artificial selection (2018–2019), followed by a period of stability despite relaxed selection pressures with the onset of the COVID-19 pandemic in 2020 (whereby populations were introduced to and maintained on heterogeneous food waste). Interestingly, despite this maintenance of genetic diversity, a sharp decline in Ne was observed in 2023–2024 across all selectively bred populations, albeit at different rates, indicative of recent genetic bottlenecks, which may have influenced population structure, possibly due to the accumulation of inbreeding, genetic drift, or ongoing natural selection pressures.

Given the observed demographic changes and the genetic differentiation among selectively bred populations, Tajima’s D analysis was performed to assess whether certain genomic regions were under selection, potentially contributing to these shifts. Tajima’s D values were calculated in non-overlapping 10,000 bp windows across all 11 populations (IFT, KR, LA, LB, LC, LD, LE, NT, SWS, SWT, and WT) to detect deviations from neutrality and identify potential selective forces.

The results revealed substantial variation in Tajima’s D values across populations and genomic regions, ranging from negative values (indicative of selective sweeps) to positive values (suggesting balancing selection), while values around zero reflected regions under neutral evolution (Figure S5A). Filtering for significantly high Tajima’s D values (>3) identified regions under strong balancing selection, with LE (48 regions) and LC (36 regions) exhibiting the highest counts among selective bred populations (LA to LE). Key genes under balancing selection included hemicentin-1 (LE, WT), aquaporin (LA), and trafficking protein subunit 6b (LE) (Supplementary file 1. “combined_annotated_final.xls”).

Conversely, selective sweeps (Tajima’s D < -3) indicated positive selection in IFT (cAMP-dependent protein kinase regulatory subunit), LA (plastin-1), and LD (uncharacterized regions), suggesting that selection may have continued shaping genetic variation even after artificial selection was relaxed.

Several genes under selection were shared across captive populations, including endoplasmic reticulum aminopeptidase 2, linked to stress adaptation (Table S3). Wild populations (WT, SWT) uniquely exhibited selection in structural organization genes, likely reflecting natural adaptation. In LA to LE, the co-occurrence of selective sweeps and balancing selection across multiple genomic regions suggested that both past artificial selection and recent demographic changes contributed to selection pressures. These patterns aligned with Figure 5, which showed recent bottlenecks in these populations.

## Discussion

This study investigated the genetic structure and differentiation of black soldier fly (BSF) populations under domestication, artificial selection, and relaxed selection, integrating mitochondrial *CO1* and genome-wide RAD-seq data. The novelty of this work lies in capturing rapid genetic shifts in selectively bred populations (Tok Wei Xian Eugene, 2019) over a short timeframe (∼5 years), within a confined laboratory environment, alongside commercial and wild-caught populations. These complementary analyses collectively illustrated the complex interplay of genetic drift, selection, and environmental adaptation in shaping population genomic landscapes.

The mitochondrial *CO1* data not only provided a foundational understanding of maternal lineages but also revealed broader patterns of dispersal and global movement. The dominance of two major haplogroups across Asia, Australia, Africa, Europe, and South America was consistent with historical introductions, likely facilitated by human-mediated commercial trade and breeding programs rather than passive range expansion (Generalovic et al., 2023). This was further supported by phylogenetic patterns showing close genetic relationships between geographically distant populations and a North American ancestor (Guilliet et al., 2022; Kaya et al., 2021). Such global movements have critical ecological implications, including potential threats to native biodiversity, maladaptation in new environments, and cases of reproductive isolation, such as those observed in South Korea (Park et al., 2017; Ståhls et al., 2020). These findings underscored the importance of maintaining accurate records of lineage origins and implementing genetic monitoring protocols in commercial and research facilities to preserve genetic integrity and optimize breeding outcomes.

### Rapid adaptation in a confined environment

While mitochondrial data hinted at large-scale patterns, genome-wide RAD-seq analysis provided a deeper exploration of population structure and dynamics. Genetic relationships revealed clear divergence between wild-caught and domesticated populations. Selectively bred lines, though originating from a common admixed founder population, showed divergence, illustrating that genetic drift and environmental adaptation continued to shape genomes even after relaxation of targeted selection (L. Hoffmann et al., 2021; Mopper & Strauss, 2013). However, this differentiation was not uniform. Some populations retained homogeneity, while others revealed unexpected levels of admixture, pointing to uneven gene flow and complex adaptive strategies even in a confined environment (Park et al., 2017).

Demographic reconstructions further highlighted how populations initially expanded under targeted artificial selection, maintained stability, and then experienced sharp declines in effective population sizes following the onset of the COVID-19 pandemic, when populations were concurrently subjected to relaxed selection on heterogeneous food waste. These recent demographic bottlenecks, likely shaped by differential adaptive strategies and breeding practices, appeared to have accelerated genetic drift and contributed to enhanced divergence among lines (Rhode et al., 2020). The interplay between these demographic changes and selection pressures was most evident in the genomic signals of selection observed across populations.

Across populations, both selective sweeps and balancing selection left distinct signatures. In selectively bred populations, balancing selection was observed in genes associated with metabolism, stress adaptation, and cellular signalling, suggesting adaptive mechanisms responding to fluctuating rearing conditions and nutritional variability (Gligorescu et al., 2023; Stajich & Hahn, 2005). Wild populations carried selection signals in genes related to structural integrity and stress resilience, consistent with environmental adaptation (Li et al., 2024; Zhan et al., 2020). The presence of shared selection signals across captive populations not only indicated common selective pressures but also suggested convergent evolution driven by similar captive environmental conditions (Lahti et al., 2009). This highlights the importance of understanding how these parallel evolutionary responses can influence breeding outcomes and long-term genetic stability. Additionally, the co-occurrence of selective sweeps and balancing selection within populations demonstrated that different parts of the genome were subjected to contrasting evolutionary forces simultaneously, with some regions losing genetic diversity due to directional selection and others maintaining diversity through fluctuating selection or heterozygote advantage, highlighting the complex dynamics of genomic evolution (Castillo & Agathos, 2019; Chantepie & Chevin, 2020; Charlesworth, 2013).

The rapid genetic divergence observed among closely related lines (LA–LE), which developed over only a few years despite relaxed selection, challenges theoretical expectations that populations revert to previous genetic states when selection ceases (Lahti et al., 2009). Instead, these findings suggest that small effective population sizes, genetic drift, and environmental pressures prevent such regression and drive continued divergence (Li et al., 2024; Rhode et al., 2020). This underscores that genomic evolution under domestication is both rapid and ongoing, with drift, demographic bottlenecks, and selective pressures reshaping population structure more quickly than anticipated, even in controlled environments (A. A. Hoffmann & Willi, 2008).

These findings emphasize the importance of understanding how different selection regimes, both intentional and unintentional, shape genetic diversity and long-term population resilience. Even under relaxed selection, genetic bottlenecks, drift, and historical selection pressures can leave lasting genomic signatures, influencing population divergence over relatively short timeframes (Lahti et al., 2009). Phenotypic differences observed in prior studies, including variation in egg production and adaptation to dietary inconsistencies, further highlight how genetic variation can drive population differentiation within a controlled environment (Gligorescu et al., 2023; Sandrock et al., 2022; Silvaraju et al., 2024; Zhang et al., 2023).

This study lays a critical foundation for future research by providing insights into the rapid evolutionary trajectory of BSF populations reared in confined settings. Building on the selective sweep analyses presented here, future transcriptomic investigations will help functionally characterize candidate genes under selection, differentiating potentially adaptive changes from neutral or deleterious variants. Together, this work contributes to the development of effective selective breeding strategies and optimized rearing practices, ensuring the genetic stability and sustainability of BSF populations in both commercial and research applications.

## Conclusion

This study demonstrated that genetic differentiation could occur rapidly in BSF populations due to their short life cycle and rapid generational turnover. Even populations derived from the same parent population exhibited genetic divergence, likely driven by differential responses to selective pressures from environmental conditions. Such differentiation may result in positive assortative mating, which, if unchecked, could reduce genetic diversity and increase the risk of population collapse. Conversely, it may also enhance adaptability, enabling certain populations to thrive under novel environmental conditions through positive selection. These findings demonstrate the dynamic relationship between genetic diversity and environmental pressures in shaping population structure, highlighting the need for biomonitoring of populations under domestication. However, further transcriptomic studies will aid in distinguishing actively expressed genes from pseudogenes, providing a clearer understanding of the impact of selection and the molecular mechanisms driving adaptation in BSF populations.

## Availability of data and materials

Supplementary figures and tables can be found in the Microsoft Word document titled “Silvaraju et al Evolution under Domestication.docx”. Analysis files and R script can be accessed via https://github.com/ReproLab/Evolution-under-domestication_BSF.git. The raw variant data (VCF file) generated from this analysis has been deposited in the European Variation Archive (EVA) under project accession PRJEB89246 and analysis accession ERZ26867197.

## References

Athanassiou, C. G., Coudron, C. L., Deruytter, D., Rumbos, C. I., Gasco, L., Gai, F., Sandrock, C., De Smet, J., Tettamanti, G., Francis, A., Petrusan, J.-I., & Smetana, S. (2024). A decade of advances in black soldier fly research: from genetics to sustainability. Journal of Insects as Food and Feed, 1–28. 10.1163/23524588-00001122

Barragan-Fonseca, K. B., Gort, G., Dicke, M., & van Loon, J. J. A. (2019). Effects of dietary protein and carbohydrate on life-history traits and body protein and fat contents of the black soldier fly Hermetia illucens. Physiological Entomology, 44(2), 148–159.

Bekker, N., Heidelbach, S., Vestergaard, S., Nielsen, M. E., Riisgaard-Jensen, M., Zeuner, E., Bahrndorff, S., & Eriksen, N. (2021). Impact of substrate moisture content on growth and metabolic performance of black soldier fly larvae. Waste Management, 127, 73–79. 10.1016/j.wasman.2021.04.028

Castillo, J. A., & Agathos, S. N. (2019). A genome-wide scan for genes under balancing selection in the plant pathogen Ralstonia solanacearum. BMC Evolutionary Biology, 19(1), 123. 10.1186/s12862-019-1456-6

Chantepie, S., & Chevin, L.-M. (2020). How does the strength of selection influence genetic correlations? Evolution Letters, 4(6), 468–478.

Charlesworth, B. (2013). Population Genetics. In S. A. Levin (Ed.), Encyclopedia of Biodiversity (Second Edition) (pp. 182–198). Academic Press. 10.1016/B978-0-12-384719-5.00116-7

Chen, G., Zhang, K., Tang, W., Li, Y., Pang, J., Yuan, X., Song, X., Jiang, L., Yu, X., Zhu, H., Wang, J., Zhang, J., & Zhang, X. (2023). Feed nutritional composition affects the intestinal microbiota and digestive enzyme activity of black soldier fly larvae. Frontiers in Microbiology, 14. https://www.frontiersin.org/journals/microbiology/articles/10.3389/fmicb.2023.1184139

Eke, M., Tougeron, K., Hamidovic, A., Tinkeu, L. S. N., Hance, T., & Renoz, F. (2023). Deciphering the functional diversity of the gut microbiota of the black soldier fly (Hermetia illucens): recent advances and future challenges. Animal Microbiome, 5(1), 40. 10.1186/s42523-023-00261-9

Gabriel Patrick. (2021, August). Top 10 black soldier fly companies feeding nutrient-rich feed to livestock. Verified Market Research.

Gebiola, M., Rodriguez, M. V, Garcia, A., Garnica, A., Tomberlin, J. K., Hopkins, F. M., & Mauck, K. E. (2023). Bokashi fermentation of brewery’s spent grains positively affects larval performance of the black soldier fly Hermetia illucens while reducing gaseous nitrogen losses. Waste Management, 171, 411–420. 10.1016/j.wasman.2023.09.033

Geller, J., Meyer, C., Parker, M., & Hawk, H. (2013). Redesign of PCR primers for mitochondrial cytochrome c oxidase subunit I for marine invertebrates and application in all-taxa biotic surveys. Molecular Ecology Resources, 13(5), 851–861. 10.1111/1755-0998.12138

Generalovic, T. N., McCarthy, S. A., Warren, I. A., Wood, J. M. D., Torrance, J., Sims, Y., Quail, M., Howe, K., Pipan, M., Durbin, R., & Jiggins, C. D. (2021). A high-quality, chromosome-level genome assembly of the Black Soldier Fly (Hermetia illucens L.). G3 Genes|Genomes|Genetics, 11(5), kab085. 10.1093/g3journal/jkab085

Generalovic, T. N., Sandrock, C., Roberts, B. J., Meier, J. I., Hauser, M., Warren, I. A., Pipan, M., Durbin, R., & Jiggins, C. D. (2023). Cryptic diversity and signatures of domestication in the Black Soldier Fly (Hermetia illucens). BioRxiv, 2023.10.21.563413. 10.1101/2023.10.21.563413

Gligorescu, A., Chen, L., Jensen, K., Moghadam, N. N., Kristensen, T. N., & Sørensen, J. G. (2023). Rapid evolutionary adaptation to diet composition in the black soldier fly (Hermetia illucens). Insects, 14(10), 821.

Greenwood, M. P., Hull, K. L., Brink-Hull, M., Lloyd, M., & Rhode, C. (2021). Feed and Host Genetics Drive Microbiome Diversity with Resultant Consequences for Production Traits in Mass-Reared Black Soldier Fly (Hermetia illucens) Larvae. Insects, 12(12). 10.3390/insects12121082

Guilliet, J., Baudouin, G., Pollet, N., & Filée, J. (2022). What complete mitochondrial genomes tell us about the evolutionary history of the black soldier fly, Hermetia illucens. BMC Ecology and Evolution, 22(1), 72. 10.1186/s12862-022-02025-6

Hoffmann, A. A., & Willi, Y. (2008). Detecting genetic responses to environmental change. Nature Reviews Genetics, 9(6), 421–432.

Hoffmann, L., Hull, K. L., Bierman, A., Badenhorst, R., Bester-Van Der Merwe, A. E., & Rhode, C. (2021). Patterns of genetic diversity and mating systems in a mass-reared black soldier fly colony. Insects, 12(6), 480.

Jiang, C., Jin, W., Tao, X., Zhang, Q., Zhu, J., Feng, S., Xu, X., Li, H., Wang, Z., & Zhang, Z. (2019). Black soldier fly larvae (Hermetia illucens) strengthen the metabolic function of food waste biodegradation by gut microbiome. Microbial Biotechnology, 12(3), 528–543.

Kaya, C., Generalovic, T. N., Ståhls, G., Hauser, M., Samayoa, A. C., Nunes-Silva, C. G., Roxburgh, H., Wohlfahrt, J., Ewusie, E. A., Kenis, M., Hanboonsong, Y., Orozco, J., Carrejo, N., Nakamura, S., Gasco, L., Rojo, S., Tanga, C. M., Meier, R., Rhode, C., … Sandrock, C. (2021). Global population genetic structure and demographic trajectories of the black soldier fly, Hermetia illucens. BMC Biology, 19(1), 94. 10.1186/s12915-021-01029-w

Keightley, P. D., Trivedi, U., Thomson, M., Oliver, F., Kumar, S., & Blaxter, M. L. (2009). Analysis of the genome sequences of three Drosophila melanogaster spontaneous mutation accumulation lines. Genome Research, 19(7), 1195–1201.

Khamis, F. M., Ombura, F. L. O., Akutse, K. S., Subramanian, S., Mohamed, S. A., Fiaboe, K. K. M., Saijuntha, W., Van Loon, J. J. A., Dicke, M., Dubois, T., Ekesi, S., & Tanga, C. M. (2020). Insights in the Global Genetics and Gut Microbiome of Black Soldier Fly, Hermetia illucens: Implications for Animal Feed Safety Control. Frontiers in Microbiology, 11. 10.3389/fmicb.2020.01538

Klammsteiner, T., Walter, A., Bogataj, T., Heussler, C. D., Stres, B., Steiner, F. M., Schlick-Steiner, B. C., Arthofer, W., & Insam, H. (2020). The Core Gut Microbiome of Black Soldier Fly (Hermetia illucens) Larvae Raised on Low-Bioburden Diets. Frontiers in Microbiology, 11. 10.3389/fmicb.2020.00993

Lahti, D. C., Johnson, N. A., Ajie, B. C., Otto, S. P., Hendry, A. P., Blumstein, D. T., Coss, R. G., Donohue, K., & Foster, S. A. (2009). Relaxed selection in the wild. Trends in Ecology & Evolution, 24(9), 487–496.

Leray, M., Yang, J. Y., Meyer, C. P., Mills, S. C., Agudelo, N., Ranwez, V., Boehm, J. T., & Machida, R. J. (2013). A new versatile primer set targeting a short fragment of the mitochondrial COI region for metabarcoding metazoan diversity: application for characterizing coral reef fish gut contents. Frontiers in Zoology, 10(1), 34. 10.1186/1742-9994-10-34

Li, X., Han, B., Liu, D., Wang, S., Wang, L., Pei, Q., Zhang, Z., Zhao, J., Huang, B., Zhang, F., Zhao, K., & Tian, D. (2024). Whole-genome resequencing to investigate the genetic diversity and mechanisms of plateau adaptation in Tibetan sheep. Journal of Animal Science and Biotechnology, 15(1), 164. 10.1186/s40104-024-01125-1

Mopper, S., & Strauss, S. Y. (2013). Genetic structure and local adaptation in natural insect populations: effects of ecology, life history, and behavior. Springer Science & Business Media.

Park, S., Choi, H., Choi, J., & Jeong, G. (2017). Population structure of the exotic black soldier fly, Hermetia illucens (Diptera: Stratiomyidae) in Korea. Korean Journal of Environment and Ecology, 31(6), 520–528.

Raimondi, S., Spampinato, G., Macavei, L. I., Lugli, L., Candeliere, F., Rossi, M., Maistrello, L., & Amaretti, A. (2020). Effect of rearing temperature on growth and microbiota composition of Hermetia illucens. Microorganisms, 8(6), 902.

Rhode, C., Badenhorst, R., Hull, K. L., Greenwood, M. P., Bester-van der Merwe, A. E., Andere, A. A., Picard, C. J., & Richards, C. (2020). Genetic and phenotypic consequences of early domestication in black soldier flies (Hermetia illucens). Animal Genetics, 51(5), 752–762. 10.1111/age.12961

Sandrock, C., Leupi, S., Wohlfahrt, J., Kaya, C., Heuel, M., Terranova, M., Blanckenhorn, W. U., Windisch, W., Kreuzer, M., & Leiber, F. (2022). Genotype-by-Diet Interactions for Larval Performance and Body Composition Traits in the Black Soldier Fly, Hermetia illucens. Insects, 13(5). 10.3390/insects13050424

Shelomi, M. (2020). Nutrient Composition of Black Soldier Fly (Hermetia illucens). In A. Adam Mariod (Ed.), African Edible Insects As Alternative Source of Food, Oil, Protein and Bioactive Components (pp. 195–212). Springer International Publishing. 10.1007/978-3-030-32952-5_13

Sheppard, D. C., Tomberlin, J. K., Joyce, J. A., Kiser, B. C., & Sumner, S. M. (2002). Rearing Methods for the Black Soldier Fly (Diptera: Stratiomyidae). Journal of Medical Entomology, 39(4), 695–698. 10.1603/0022-2585-39.4.695

Silvaraju, S., Zhang, Q., Kittelmann, S., & Puniamoorthy, N. (2024). Genetics, age, and diet influence gut bacterial communities and performance of black soldier fly larvae (Hermetia illucens). 10.21203/rs.3.rs-4661186/v1

Slagboom, M., Nielsen, H. M., Kargo, M., Henryon, M., & Hansen, L. S. (2024). The effect of phenotyping, adult selection, and mating strategies on genetic gain and rate of inbreeding in black soldier fly breeding programs. Genetics Selection Evolution, 56(1), 71. 10.1186/s12711-024-00938-y

Spranghers, T., Ottoboni, M., Klootwijk, C., Ovyn, A., Deboosere, S., De Meulenaer, B., Michiels, J., Eeckhout, M., De Clercq, P., & De Smet, S. (2017). Nutritional composition of black soldier fly (Hermetia illucens) prepupae reared on different organic waste substrates. Journal of the Science of Food and Agriculture, 97(8), 2594–2600. 10.1002/jsfa.8081

Ståhls, G., Meier, R., Sandrock, C., Hauser, M., Šašic Zoric, L., Laiho, E., Aracil, A., Doderović, J., Badenhorst, R., Unadirekkul, P., Mohd Adom, N. A. B., Wein, L., Richards, C., Tomberlin, J. K., Rojo, S., Veselić, S., & Parviainen, T. (2020). The puzzling mitochondrial phylogeography of the black soldier fly (Hermetia illucens), the commercially most important insect protein species. BMC Evolutionary Biology, 20(1), 60. 10.1186/s12862-020-01627-2

Stajich, J. E., & Hahn, M. W. (2005). Disentangling the Effects of Demography and Selection in Human History. Molecular Biology and Evolution, 22(1), 63–73. 10.1093/molbev/msh252

Tok Wei Xian Eugene, A. (2019). Title IMPROVING BLACK SOLDIER FLIES FOR FOOD WASTE RECYCLING THROUGH ARTIFICIAL SELECTION. https://scholarbank.nus.edu.sg/handle/10635/165828

Vandeweyer, D., Bruno, D., Bonelli, M., IJdema, F., Lievens, B., Crauwels, S., Casartelli, M., Tettamanti, G., & De Smet, J. (2023). Bacterial biota composition in gut regions of black soldier fly larvae reared on industrial residual streams: revealing community dynamics along its intestinal tract. Frontiers in Microbiology, 14. https://www.frontiersin.org/journals/microbiology/articles/10.3389/fmicb.2023.1276187

Yakti, W., Schulz, S., Marten, V., Mewis, I., Padmanabha, M., Hempel, A. J., Kobelski, A., Streif, S., & Ulrichs, C. (2022). The Effect of Rearing Scale and Density on the Growth and Nutrient Composition of Hermetia illucens (L.) (Diptera: Stratiomyidae) Larvae. Sustainability (Switzerland), 14(3). 10.3390/su14031772

Yang, F., Tomberlin, J. K., & Jordan, H. R. (2021). Starvation Alters Gut Microbiome in Black Soldier Fly (Diptera: Stratiomyidae) Larvae. Frontiers in Microbiology, 12. 10.3389/fmicb.2021.601253

Zhan, S., Fang, G., Cai, M., Kou, Z., Xu, J., Cao, Y., Bai, L., Zhang, Y., Jiang, Y., Luo, X., Xu, J., Xu, X., Zheng, L., Yu, Z., Yang, H., Zhang, Z., Wang, S., Tomberlin, J. K., Zhang, J., & Huang, Y. (2020). Genomic landscape and genetic manipulation of the black soldier fly Hermetia illucens, a natural waste recycler. Cell Research, 30(1), 50–60. 10.1038/s41422-019-0252-6

Zhang, Q.-H., Silvaraju, S., Unadirekkul, P., Lim, N. W., Heng, C. W., Liu, M. H., & Puniamoorthy, N. (2023). Laboratory-adapted and wild-type black soldier flies express differential plasticity in bioconversion and nutrition when reared on urban food waste streams. Journal of the Science of Food and Agriculture, n/a(n/a). 10.1002/jsfa.13039

Zhineng, Y., Ying, M., Bingjie, T., Rouxian, Z., & Qiang, Z. (2021). Intestinal microbiota and functional characteristics of black soldier fly larvae (Hermetia illucens). Annals of Microbiology, 71(1), 13. 10.1186/s13213-021-01626-8

Ebeneezar, S., D., L. P., C.s., T., N.s., J., R., S., S., C., P., S., & P., V. (2021). Nutritional evaluation, bioconversion performance and phylogenetic assessment of black soldier fly (Hermetia illucens, Linn. 1758) larvae valorized from food waste. Environmental Technology & Innovation, 23, 101783. 10.1016/j.eti.2021.101783

Srivathsan, A., Baloğlu, B., Wang, W., Tan, W. X., Bertrand, D., Ng, A. H. Q., Boey, E. J. H., Koh, J. J. Y., Nagarajan, N., & Meier, R. (2018). A MinIONTM-based pipeline for fast and cost-effective DNA barcoding. Molecular Ecology Resources, 18(5), 1035–1049. 10.1111/1755-0998.12890

Srivathsan, A., Lee, L., Katoh, K., Hartop, E., Kutty, S. N., Wong, J., Yeo, D., & Meier, R. (2021). ONTbarcoder and MinION barcodes aid biodiversity discovery and identification by everyone, for everyone. BMC Biology, 19(1), 217. 10.1186/s12915-021-01141-x

