## Supplementary data for "Evolution under Domestication: Genetic differentiation in black soldier fly (*Hermetia illucens*) populations subjected to recent selective breeding"

Table S1: Table showing the number of BSF sequences adopted from various publications for analyses in the present study, and their GenBank Accession Numbers.

| **Title** | **Year of publication** | **GenBank Accession Numbers** | **Number of sequences yielded** |
|---|---|---|---|
| The puzzling mitochondrial phylogeography of the black soldier fly (Hermetia illucens), the commercially most important insect protein species | 2020 | MT151287, MT151288, LR778156, LR778157, MT178507, MT181121, LR778158, LR792223, LR778159, LR778160, JQ344949, MT178504, MT178505, LR778161, LR778162, LR792224, MT178509, MT186669, MT178484, MT178485, MT178486, MT178501, MT178510, MT178508, MT178500, LR792225, LR792227, LR792226, LR778207, MT181123, MT178491, MT178492, MT178488, MT178458, MT178459, MT178460, MT178461, MT181122, LR792228, LR792229, LR792230, LR792231, LR792232, LR778212, LR778213, LR792234, LR778211, LR778209, LR778210, LR792233, LR778208, LR792235, LR778205, LR778206, LR792236, LR792237, LR778204, LR792238, LR778203, LR778195, LR778202, LR778197, LR778193, LR778198, LR778199, LR778196, LR778194, LR778192, LR778201, LR778200, LR792239, LR792240, LR792241, LR778154, LR778155 | 75 |
| Population Structure of the Exotic Black Soldier Fly, Hermetia illucens (Diptera: Stratiomyidae) in Korea | 2018 | FJ794325~FJ794425, HQ541184 ~HQ541321, and KF500241~KF500278 | 244 |
| Nutritional evaluation, bioconversion performance and phylogenetic assessment of black soldier fly (Hermetia illucens, Linn. 1758) larvae valorized from food waste | 2021 | MW173681, MW278893–MW278894, MZ148804–MZ148806. | 6 |
| Genotype-by-Diet Interactions for Larval Performance and Body Composition Traits in the Black Soldier Fly, Hermetia illucens | 2022 | LR792261, LR792262, LR792223, LR792267 | 4 |

Table S2. Summary of mitochondrial *CO1* haplotype diversity (*Hd*), nucleotide diversity ($\pi$), and overall genetic diversity indices across BSF populations.

| Population | Hd | π | Diversity |
|---|---|---|---|
| LA | 0 | 0 | Nil |
| LB | 0.521 | 0.002 | Moderate Hd; Low π |
| LC | 0.189 | 0.0006 | Low |
| LD | 0 | 0 | Nil |

| | | | |
|---|---|---|---|
| LE | 0 | 0 | Nil |
| WT | 0 | 0 | Nil |
| SWS | 0 | 0 | Nil |
| SWT | 0.268 | 0.0009 | Low |
| KR | 0.4 | 0.018 | Low Hd; Moderate π |
| IFT | 0.419 | 0.021 | Low Hd; High π |
| NT | 0.762 | 0.025 | High |

Table S3. List of shared genes across BSF populations, including gene identifiers, populations sharing those genes, and their respective annotated functions.

| Gene | Shared Populations | Function |
|---|---|---|
| LOC119653827 | LC, LE, SWT | 1-acyl-sn-glycerol-3-phosphate acyltransferase beta,2C transcript variant X2 |
| LOC119656787 | LB, SWT | anoctamin-4,2C transcript variant X3 |
| LOC119650970 | IFT, WT | beta-1,2C4-glucuronyltransferase 1,2C transcript variant X3 |
| LOC119660900 | LC, LB | bromodomain-containing protein DDB_G0280777-like,2C transcript variant X1 |
| LOC119656007 | LB, IFT, SWS, LE, SWT | cAMP-dependent protein kinase type II regulatory subunit,2C transcript variant X1 |
| LOC119646149 | LC, LD, IFT, LA, SWS, WT, LE, LB, SWT | cGMP-dependent protein kinase,2C isozyme 2 forms cD4/T1/T3A/T3B,2C transcript variant X9 |
| LOC119650441 | SWT, WT | coiled-coil domain-containing protein 102A,2C transcript variant X3 |
| LOC119652886 | IFT, LB, LE | E3 ubiquitin-protein ligase RNF4-like,2C transcript variant X2 |
| LOC119646603 | IFT, LB, LC, NT, SWS, SWT, LA, LD, LE, WT | endoplasmic reticulum aminopeptidase 2,2C transcript variant X5 |
| LOC119650309 | IFT, LA, LC, SWT | facilitated trehalose transporter Tret1,2C transcript variant X10 |
| LOC119650467 | WT, IFT, SWT | FH1/FH2 domain-containing protein 3,2C transcript variant X7 |
| LOC119647100 | IFT, LC, SWT | filamin-A,2C transcript variant X7 |
| LOC119647447 | LA, LB, LC, LE, WT | four and a half LIM domains protein 2,2C transcript variant X1 |
| LOC119661033 | LD, LE, WT | GIGYF family protein CG11148,2C transcript variant X3 |
| LOC119651895 | LD, LE, WT | GTP-binding protein 2,2C transcript variant X2 |
| LOC119649806 | LA, LC, LD, LE, SWS, WT | hemicentin-1,2C transcript variant X15 |
| LOC119649052 | KR, LA, LC, LE, WT | inactive dipeptidyl peptidase 10,2C transcript variant X1 |
| LOC119651953 | NT, SWS | ion transport peptide,2C transcript variant X4 |
| LOC119656907 | KR, LE, SWS | kelch-like protein 30,2C transcript variant X4 |
| LOC119653720 | LC, SWT | mucin-17,2C transcript variant X7 |
| LOC119653731 | LA, LC, LD | NAD kinase-like,2C transcript variant X2 |
| LOC119647833 | LA, LC, LD, SWS, SWT | neural-cadherin,2C transcript variant X10 |
| LOC119650737 | SWT, SWS | nucleolysin TIAR-like,2C transcript variant X4 |

| | | |
|---|---|---|
| LOC119648386 | LC, LD, NT, IFT, WT, LB | potassium voltage-gated channel subfamily KQT member 1-like,2C transcript variant X2 |
| LOC119651360 | IFT, LE | probable cytochrome P450 9f2,2C transcript variant X2 |
| LOC119652389 | LA, LB, LC, LE | probable Dol-P-Man:Man(7)GlcNAc(2)-PP-Dol alpha-1,2C6-mannosyltransferase,2C transcript variant X2 |
| LOC119648147 | LA, LC | protein SCAI,2C transcript variant X10 |
| LOC119653451 | LB, LD, LE, NT, SWT, LA | putative inorganic phosphate cotransporter,2C transcript variant X1 |
| LOC119655696 | SWT, LB, LC, LD, LE | ras GTPase-activating protein raskol,2C transcript variant X1 |
| LOC119650713 | LC, LE, LA, LB | RNA pseudouridylate synthase domain-containing protein 2,2C transcript variant X1 |
| LOC119653580 | IFT, NT | serine/threonine-protein kinase grp,2C transcript variant X1 |
| LOC119653263 | LA, LD | serine/threonine-protein kinase svkA,2C transcript variant X4 |
| LOC119651411 | LA, WT | set1/Ash2 histone methyltransferase complex subunit ASH2,2C transcript variant X2 |
| LOC119647268 | SWT, LA, WT | SRSF protein kinase 3,2C transcript variant X2 |
| LOC119659823 | IFT, LC | trypsin-3-like,2C transcript variant X1 |
| LOC119647408 | LA, LB, LC | tumor protein p53-inducible nuclear protein 1,2C transcript variant X2 |
| 893 Genes | Up to all populations | Unknown |

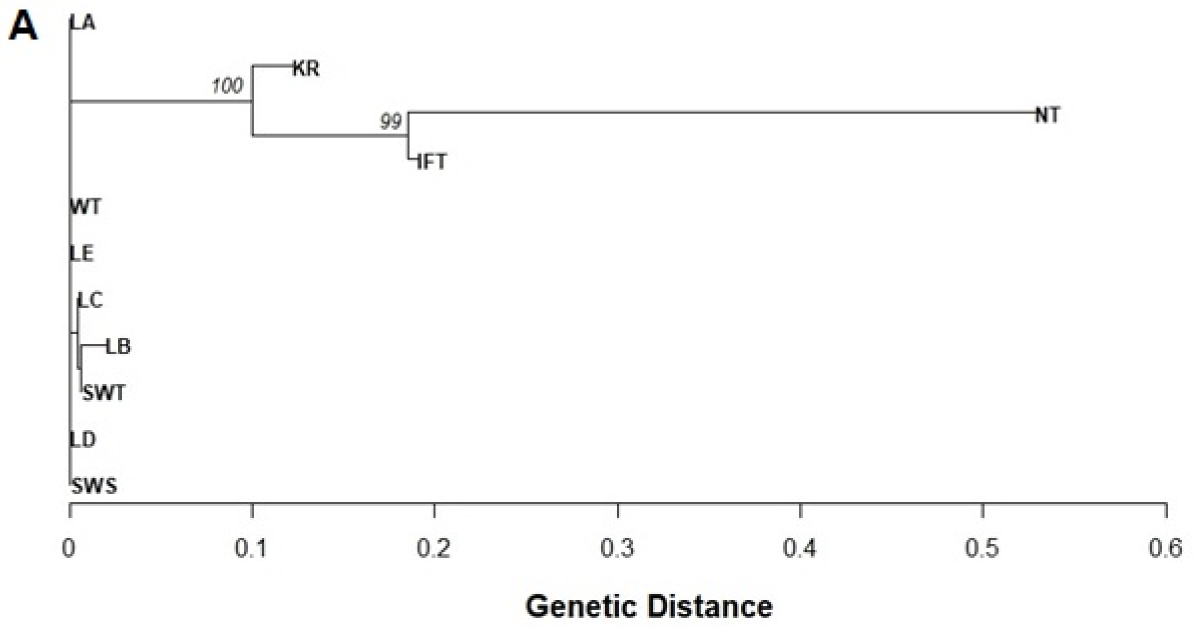

B

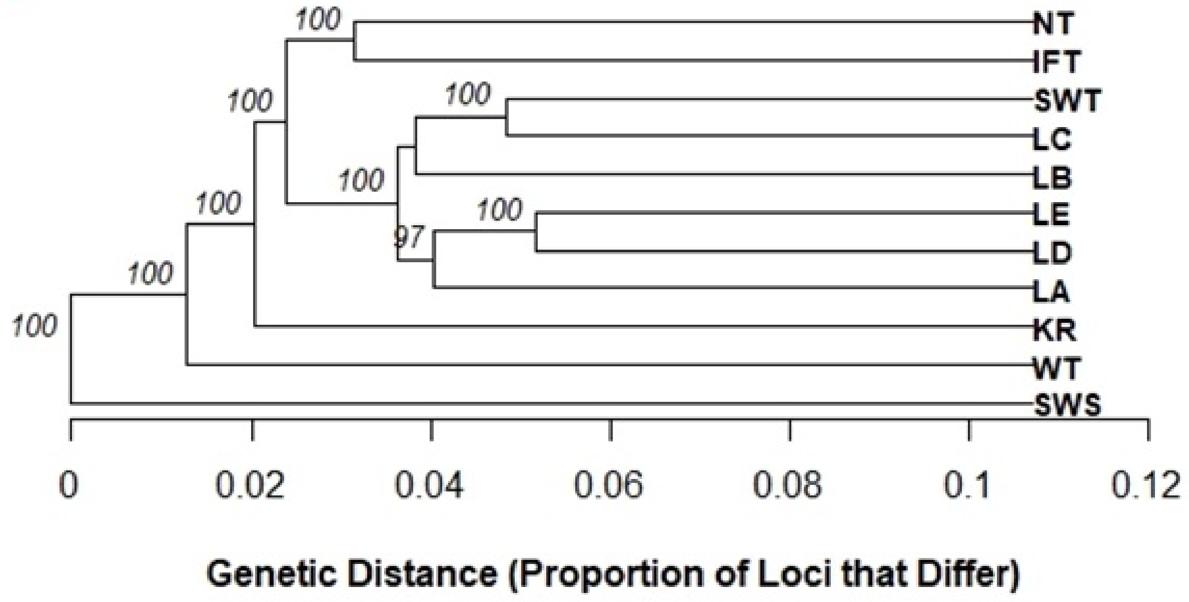

Figure S1. A) Neighbour-Joining phylogenetic tree illustrating the genetic distance among samples from genetic populations used in this study. Bootstrap values (%) at each node represent the support for the branching pattern, with values above 95% indicating strong support. B) Phylogenetic tree illustrating the genetic distance among populations used in this study. The tree was constructed using UPGMA (Unweighted Pair Group Method with Arithmetic Mean) based on the proportion of loci that differ between samples. Bootstrap values (%) at each node represent the support for the branching pattern, with values above 95% indicating strong support.

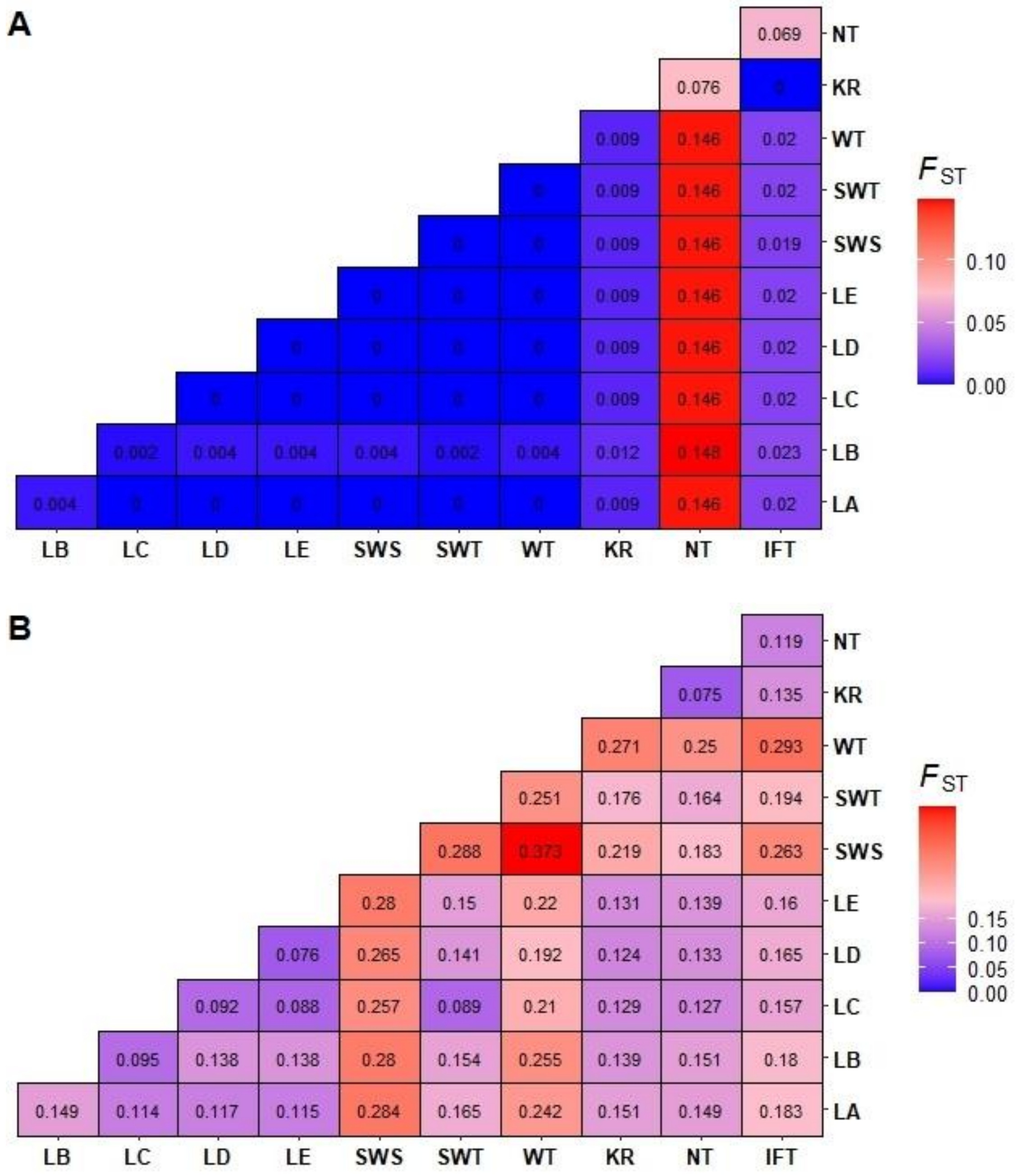

Figure S2. (A) Heatmap illustrating the Fst values based on mitochondrial *CO1* data measuring genetic differentiation among the different BSF populations. The colours indicate the degree of differentiation between populations ranging from 0 (blue; no genetic differentiation), to 0.146 (red; moderate genetic differentiation). (B) Heatmap illustrating the Fst values based on RAD-seq data (SNPs). The colours indicate the degree of differentiation ranging from 0 (blue; no genetic differentiation), to 0.373 (red; high genetic differentiation).

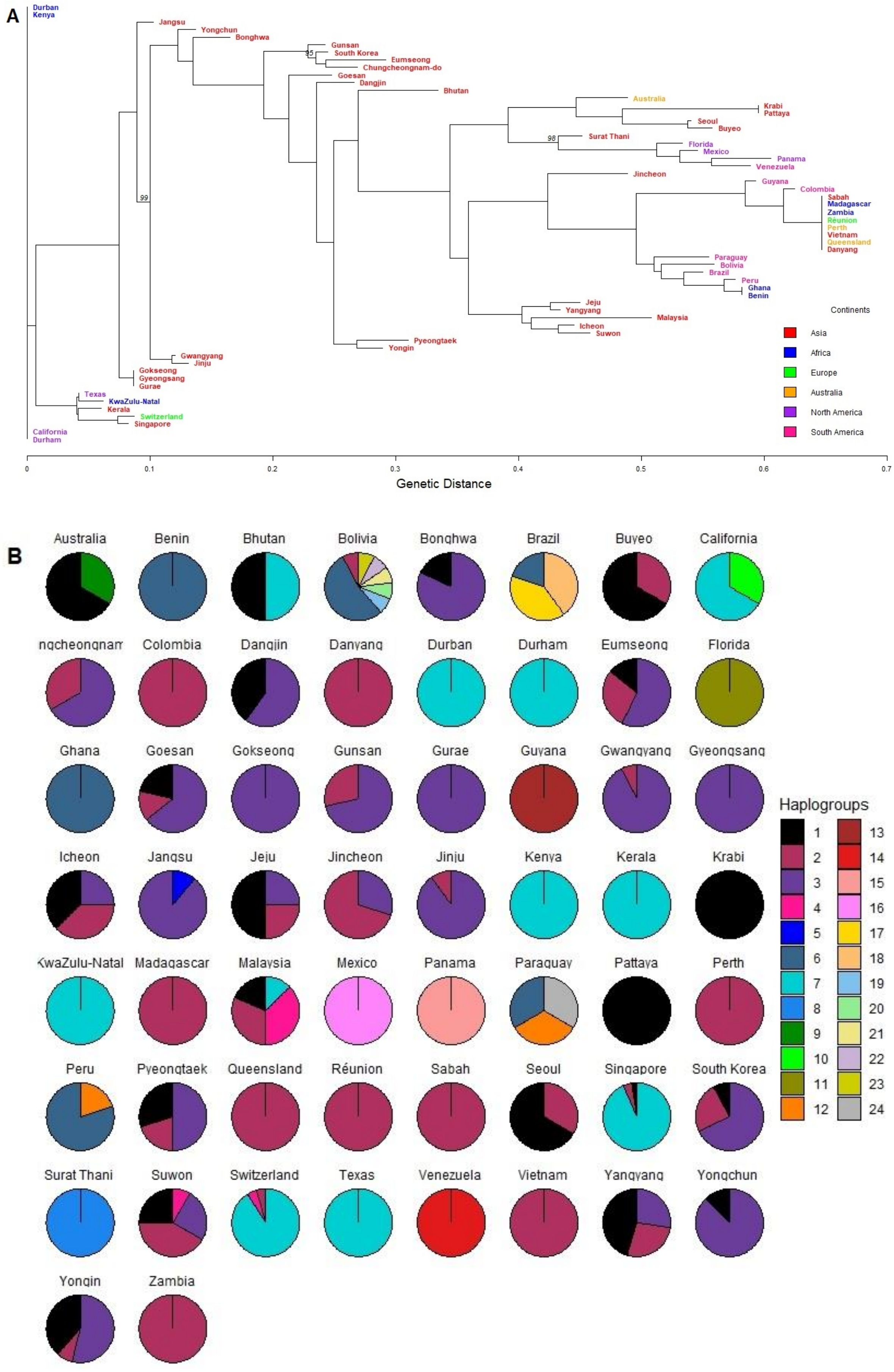
A
Durban
Kenya
Jangsu
Yongchun
Bonghwa
Gunsan
South Korea
Eumseong
Chungcheongnam-do
Goesan
Dangjin
Bhutan
Australia
Krabi
Pattaya
Seoul
Buyeo
Surat Thani
Florida
Mexico
Panama
Venezuela
Jincheon
Guyana
Colombia
Sabah
Madagascar
Zambia
Réunion
Perth
Vietnam
Queensland
Danyang
Paraguay
Bolivia
Brazil
Peru
Ghana
Benin
Jeju
Yangyang
Malaysia
Icheon
Suwon
Pyeongtaek
Yongin
Gwangyang
Jinju
Gokseong
Gyeongsang
Gurae
Texas
KwaZulu-Natal
Kerala
Switzerland
Singapore
California
Durham
95
98
99
Continents
Asia
Africa
Europe
Australia
North America
South America
0
0.1
0.2
0.3
0.4
0.5
0.6
0.7
Genetic Distance
B
Australia
Benin
Bhutan
Bolivia
Bonghwa
Brazil
Buyeo
California
ngcheongnam
Colombia
Dangjin
Danyang
Durban
Durham
Eumseong
Florida
Ghana
Goesan
Gokseong
Gunsan
Gurae
Guyana
Gwangyang
Gyeongsang
Icheon
Jangsu
Jeju
Jincheon
Jinju
Kenya
Kerala
Krabi
KwaZulu-Natal
Madagascar
Malaysia
Mexico
Panama
Paraguay
Pattaya
Perth
Peru
Pyeongtaek
Queensland
Réunion
Sabah
Seoul
Singapore
South Korea
Surat Thani
Suwon
Switzerland
Texas
Venezuela
Vietnam
Yangyang
Yongchun
Yongin
Zambia
Haplogroups

Figure S3. (A) Neighbour-Joining phylogenetic tree illustrating the genetic distance among samples from global *CO1* dataset publicly available. Each region or origin is colour coded corresponding to respective continents. Bootstrap values (%) at each node represent the support for the branching pattern, with values above 90% indicating strong support. (B) Pie charts illustrating haplogroups based on global *CO1* dataset publicly available. Each pie chart represents a specific region and is divided into segments corresponding to haplogroup frequencies. The colours represent different haplogroups (1–24) as shown in the legend, which were assigned based on a hierarchical clustering algorithm applied to genetic distance data (h = 2) grouping samples with up to 2 base pair (bp) differences into the same haplogroup.

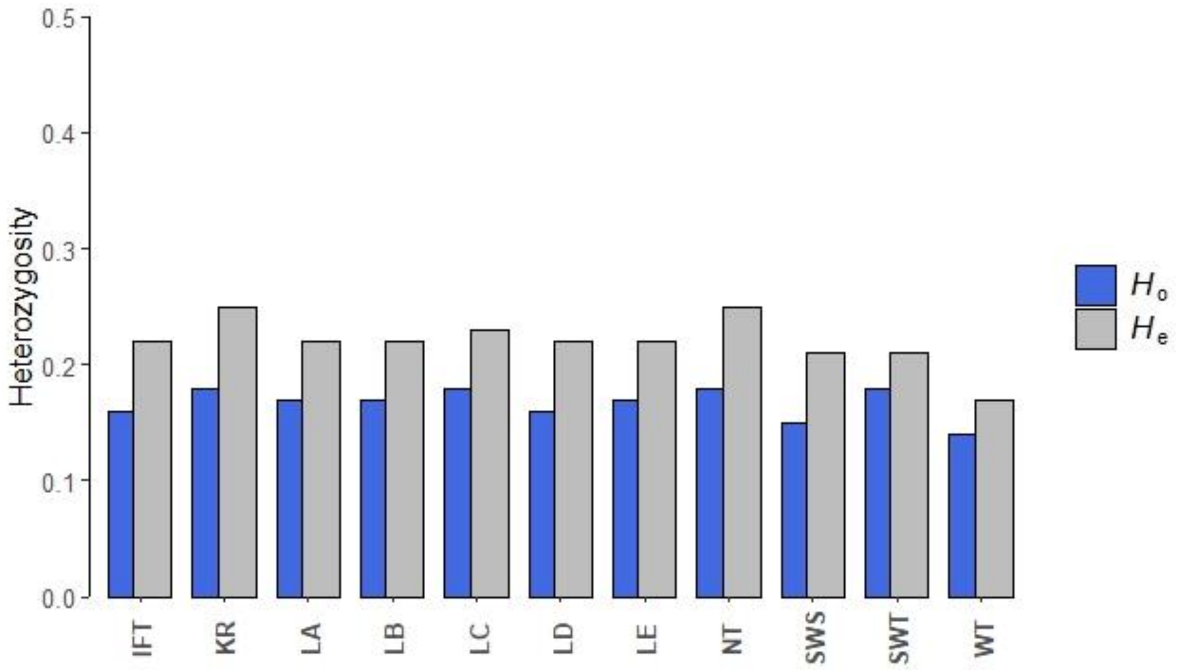

Figure S4. Observed ($H_o$, blue) and expected ($H_e$, gray) heterozygosity across BSF populations based on RAD-seq SNP data. Expected heterozygosity ($H_e$) reflects the genetic diversity predicted under Hardy-Weinberg equilibrium, while observed heterozygosity ($H_o$) represents the actual genetic diversity detected in the populations.

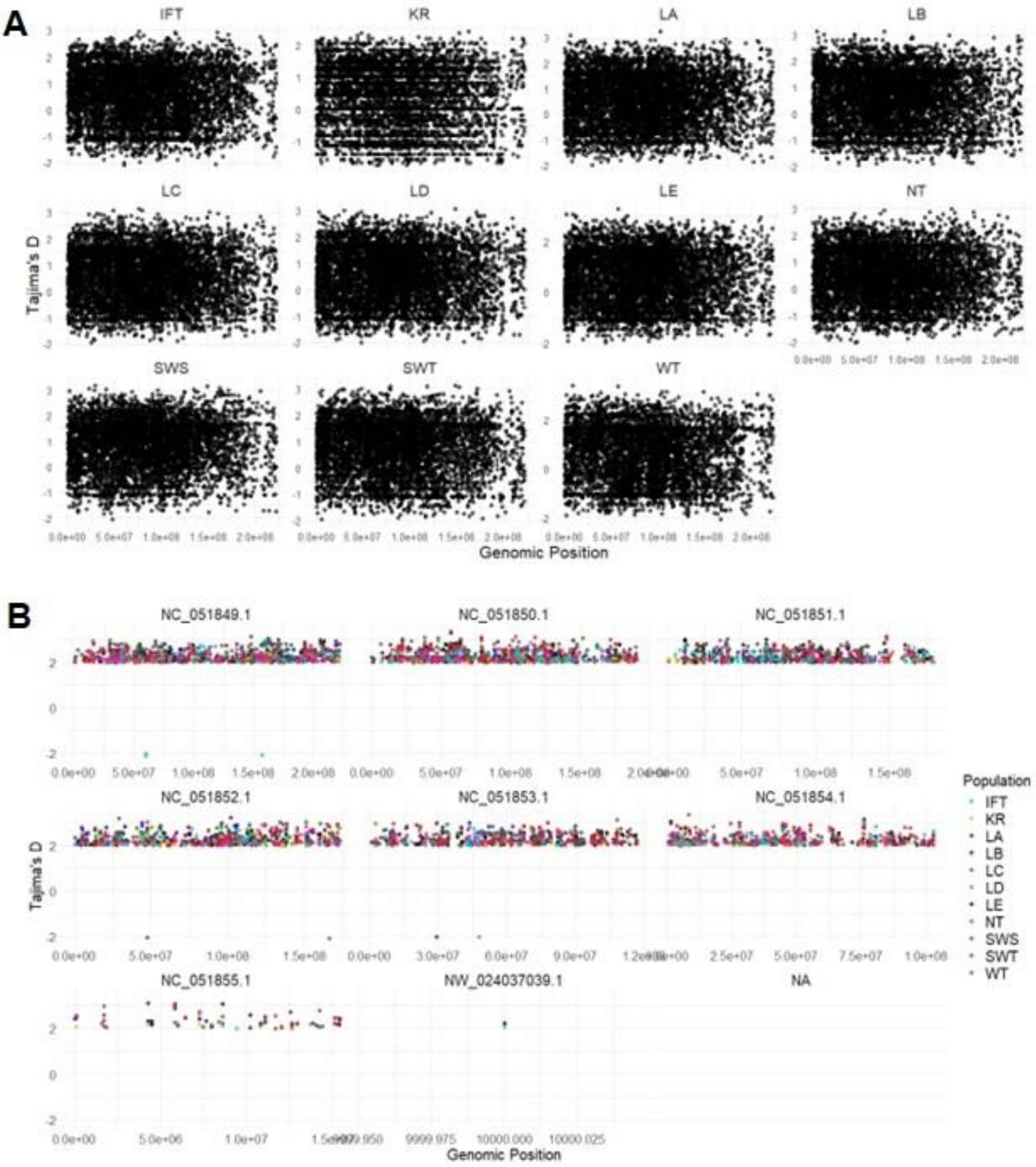
A
IFT
KR
LA
LB
LC
LD
LE
NT
SWS
SWT
WT
Tajima's D
Genomic Position
B
NC_051849.1
NC_051850.1
NC_051851.1
NC_051852.1
NC_051853.1
NC_051854.1
NC_051855.1
NW_024037039.1
NA
Tajima's D
Genomic Position
Population
IFT
KR
LA
LB
LC
LD
LE
NT
SWS
SWT
WT

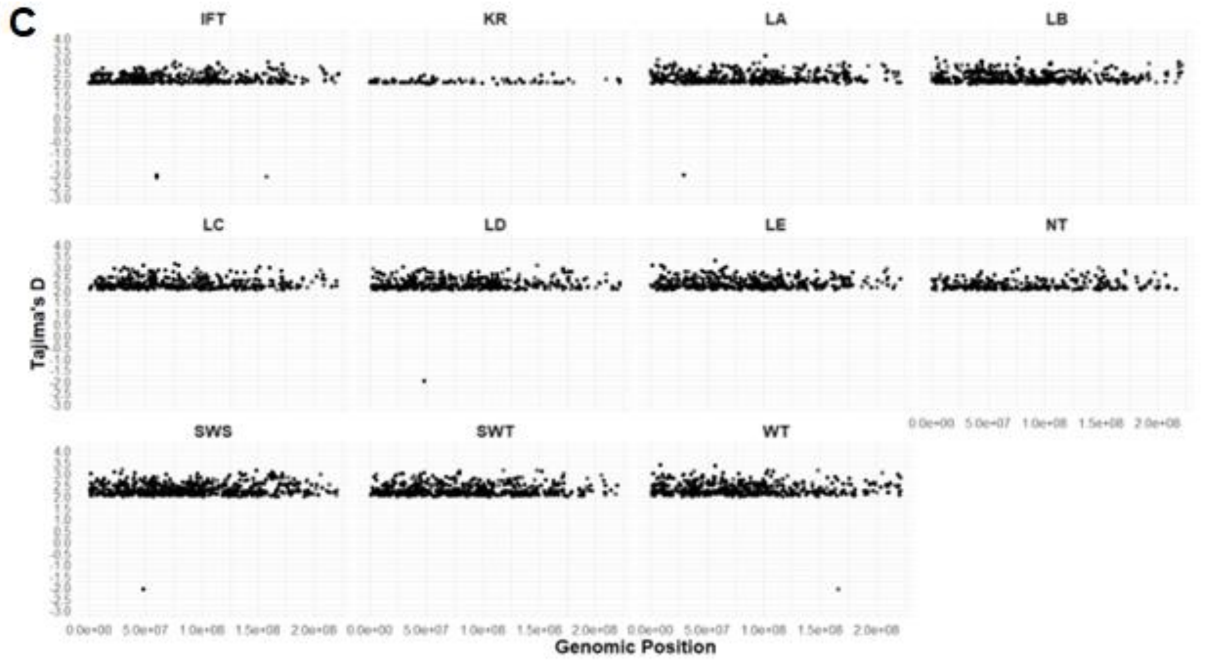

Figure S5. (A) Genome-wide distribution of Tajima's D values across different BSF populations. The values are plotted across genomic positions, with higher Tajima's D values suggesting balancing selection or population contraction, while lower values may indicate recent selective sweeps or population expansion. Values near zero suggest neutral evolution, where genetic variation aligns with expectations under mutation-drift equilibrium. B) Genome-wide distribution of Tajima's D values across the studied BSF populations. Only genomic positions with Tajima's D scores of Tajima's D > 2 (balancing selection or population contraction) and Tajima's D < -2 (selective sweep or population expansion) were plotted. C) Chromosome-level distribution of Tajima's D values across significant genomic regions in BSF populations after filtering (Tajima's D > 2 for balancing selection and < -2 for selective sweeps). Each panel represents a distinct chromosome, with coloured points indicating Tajima's D values for respective populations.
